# Deep learning enables genetic analysis of the human thoracic aorta

**DOI:** 10.1101/2020.05.12.091934

**Authors:** James P. Pirruccello, Mark D. Chaffin, Stephen J. Fleming, Alessandro Arduini, Honghuang Lin, Shaan Khurshid, Elizabeth L. Chou, Samuel N. Friedman, Alexander G. Bick, Lu-Chen Weng, Seung Hoan Choi, Amer-Denis Akkad, Puneet Batra, Nathan R. Tucker, Amelia W. Hall, Carolina Roselli, Emelia J. Benjamin, Shamsudheen K. Vellarikkal, Rajat M. Gupta, Christian M. Stegman, Jennifer E. Ho, Udo Hoffmann, Steven A. Lubitz, Anthony Philippakis, Mark E. Lindsay, Patrick T. Ellinor

## Abstract

The aorta is the largest blood vessel in the body, and enlargement or aneurysm of the aorta can predispose to dissection, an important cause of sudden death. While rare syndromes have been identified that predispose to aortic aneurysm, the common genetic basis for the size of the aorta remains largely unknown. By leveraging a deep learning architecture that was originally developed to recognize natural images, we trained a model to evaluate the dimensions of the ascending and descending thoracic aorta in cardiac magnetic resonance imaging. After manual annotation of just 116 samples, we applied this model to 3,840,140 images from the UK Biobank. We then conducted a genome-wide association study in 33,420 individuals, revealing 68 loci associated with ascending and 35 with descending thoracic aortic diameter, of which 10 loci overlapped. Integration of common variation with transcriptome-wide analyses, rare-variant burden tests, and single nucleus RNA sequencing prioritized *SVIL*, a gene highly expressed in vascular smooth muscle, that was significantly associated with the diameter of the ascending and descending aorta. A polygenic score for ascending aortic diameter was associated with a diagnosis of thoracic aortic aneurysm in the remaining 391,251 UK Biobank participants who did not undergo imaging (HR = 1.44 per standard deviation; P = 3.7·10^−12^). Defining the genetic basis of the diameter of the aorta may enable the identification of asymptomatic individuals at risk for aneurysm or dissection and facilitate the prioritization of potential therapeutic targets for the prevention or treatment of aortic aneurysm. Finally, our results illustrate the potential for rapidly defining novel quantitative traits derived from a deep learning model, an approach that can be more broadly applied to biomedical imaging data.

---

Aortic aneurysm, a pathologic enlargement of the aorta, is common, having a prevalence of approximately ∼1% of people in industrialized nations^1^. Over time, the enlarged aorta progressively expands; this process can lead to aortic dissection and rupture, which are the most catastrophic complications of aortic aneurysm and important causes of sudden cardiac death. Currently, the most effective preventive therapy is surgical repair of the aorta, a morbid operation that is only performed when aneurysms are detected prior to aortic dissection. However, timely detection is uncommon because thoracic aortic aneurysm is typically asymptomatic until the time of dissection or rupture. Unlike abdominal aortic aneurysm which has clinical screening guidelines, population screening for thoracic aortic aneurysm is not routinely performed^2,3^.

Consequently, the epidemiological and genetic contributions to aortic aneurysm have long been of interest to investigators. Clinical studies have suggested the close association of aneurysms of the descending thoracic aorta with atherosclerosis and lifestyle associated risk factors, while those of the ascending aorta occur in younger patients, sometimes associated with pathogenic genetic predisposition^4–6^. Mutations in several genes have been associated with ascending aortic aneurysms, but the small number of implicated genes is mostly limited to highly penetrant Mendelian loci identified in family studies^7–9^. Thus, there is an urgent need to identify the genetic basis for variation in aortic size in order to enable the development of new therapeutic targets for medical intervention and to identify at-risk individuals with aortic aneurysms.

We hypothesized that the size of the thoracic aorta is a complex trait, with contributions from common genetic variants. However, as the ascending and descending thoracic aorta have not only separate biological origins^10,11^, but also separate clinical risk factors^12^, we chose to quantify these aortic regions independently. Therefore, we used deep learning to localize and measure the ascending and descending thoracic aorta in 37,910 UK Biobank participants who have undergone cardiac magnetic resonance imaging (MRI) (**Table 1**). By retraining pre-existing models developed for a different purpose (recognition of objects in common images), we were able to extract data from all 3,840,140 images in the dataset after manually annotating only 116 images^13,14^. Specifically, we performed semantic segmentation—the task of identifying and labeling all pixels that comprise an object in an image—on the cross-sectional images of the ascending and descending thoracic aorta. To achieve this, we used a deep convolutional neural network that was designed with a U-Net architecture^13,14^. Such an architecture is designed to permit a model to recognize both the semantic content of an input (such as the presence of the aorta), and the fine-grained localization of that semantic label within the input image. This model used an encoder that had been pre-trained on ImageNet, which is a natural-image classification dataset; therefore, instead of starting with random weights, the model was initialized with weights that are helpful for processing images, reducing the amount of manual annotation and model training necessary to achieve good results. To recognize the aorta, this pre-trained model was retrained using only 92 manually annotated cardiac MRI still-frame images, achieving 97.4% pixel categorization accuracy in a held-out validation set of 24 additional manually annotated images. The deep learning model was then applied to all 3,840,140 available images (**Figure 1**). Quality control was performed to remove images in which the aorta was deemed to be incorrectly recognized according to one or more heuristics (see **Online Methods** and the sample flow diagram in **Supplementary Figure 1**).

**Table 1.** Baseline characteristics of participants

|  | Women | Men |
| --- | --- | --- |
| N | 17516 | 15904 |
| Age at time of MRI | 63.5 (7.40) | 64.7 (7.6) |
| BMI (kg/m<sup>2</sup>) | 25.8 (4.3) | 27.0 (3.6) |
| Height (cm) | 163 (6.2) | 177 (6.6) |
| Weight (kg) | 68.9 (12.1) | 84.1 (12.6) |
| Systolic blood pressure (mmHg) | 131 (18) | 139 (16) |
| Diastolic blood pressure (mmHg) | 79 (10) | 84 (10) |
| American standard drinks per week | 4.9 (5.5) | 6.1 (7.0) |
| Smoking status |  |  |
| Current | 877 (5 %) | 1163 (7 %) |
| Never | 11243 (64 %) | 9071 (57 %) |
| Prefer not to answer | 32 (0 %) | 31 (0 %) |
| Previous | 5360 (31 %) | 5636 (35 %) |
| Unknown | 4 (0 %) | 3 (0 %) |
| Pack years of smoking | 3.6 (9.0) | 5.7 (12.7) |
| History of aortic procedures | 0 (0 %) | 0 (0 %) |
| Ascending aorta, minor axis length (cm) | 2.72 (0.30) | 3.00 (0.33) |
| Descending aorta, minor axis length (cm) | 1.93 (0.17) | 2.19 (0.19) |

**Figure 1:**
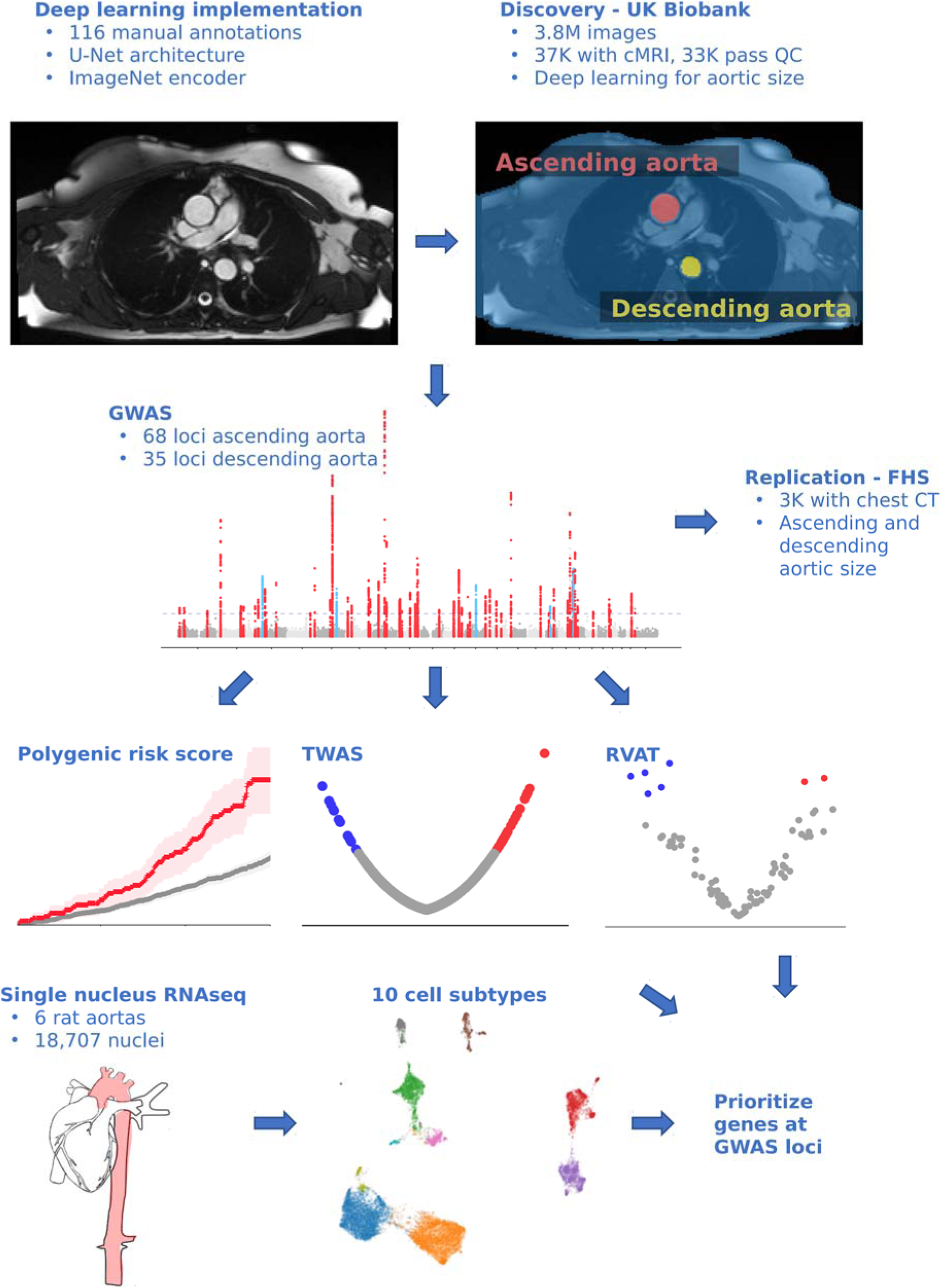
Study overview.

Having identified which pixels represent the aorta, we were able to determine the length of the minor axis (i.e., the diameter) of the ascending and descending thoracic aorta at their maximum size during the cardiac cycle (with descriptive statistics available in **Supplementary Table 1** and **Supplementary Figure 2**) and treated these as our primary phenotypes for subsequent analyses. We characterized the relationship between the aortic diameter and other anthropometric measurements and diseases in the UK Biobank (**Supplementary Note**; **Supplementary Tables 2-3**; **Supplementary Figure 3**).

We next sought to understand the common genetic basis for variation in the size of the ascending and descending thoracic aorta. We confirmed that both traits were highly heritable: the single nucleotide polymorphism (SNP) heritability of the size of the ascending aorta was 61% (95% CI 58%-65%), while that of the descending aorta was 49% (95% CI 46%-53%). We then conducted a genome-wide association study (GWAS), testing 16,563,893 imputed variants with minor allele frequency (MAF) > 0.001 for association with these phenotypes in 33,420 participants from the UK Biobank.

We identified 68 independent loci associated with the diameter of the ascending aorta at a commonly used genome-wide significance threshold (P < 5×10^−8^)(**Table 2, Figure 2A and 2B**). Of these, 64 loci were novel, and one was found on the X chromosome. In the descending aorta, we identified 35 genome-wide significant loci of which 32 were novel and one was located on the X chromosome. In total, we identified 93 loci, of which 10 were associated at genome-wide significance with both traits (**Figure 2C**). Inflation was well controlled (**Supplementary Table 4**), and no autosomal lead SNP deviated from Hardy-Weinberg Equilibrium (HWE) with P < 1×10^−6^.

**Table 2.** GWAS Loci

Ascending thoracic aorta
| SNP | CHR | BP | Effect Allele | Other Allele | EAF | INFO | BETA | P Value | Nearest Gene | Prior |
| --- | --- | --- | --- | --- | --- | --- | --- | --- | --- | --- |
| rs2871651 | 1 | 9434969 | C | T | 0.58 | 0.99 | -0.047 | 1.30E-09 | SPSB1 |  |
| rs61776719 | 1 | 38461319 | C | A | 0.45 | 1.00 | 0.043 | 1.40E-09 | SF3A3 |  |
| rs3768274 | 1 | 41951383 | C | T | 0.50 | 0.98 | -0.040 | 1.70E-08 | EDN2 |  |
| rs41519044 | 1 | 185694813 | T | A | 0.73 | 0.99 | -0.050 | 8.90E-09 | HMCN1 |  |
| rs10174214 | 2 | 19723613 | A | C | 0.32 | 1.00 | 0.090 | 6.00E-34 | OSR1 |  |
| rs2381688 | 2 | 145838542 | A | G | 0.35 | 0.97 | -0.041 | 1.40E-08 | ZEB2 |  |
| rs10186643 | 2 | 148803587 | G | T | 0.68 | 0.99 | -0.050 | 4.10E-10 | MBD5 |  |
| rs35930173 | 2 | 164924332 | G | A | 0.76 | 0.98 | -0.049 | 5.80E-10 | FIGN |  |
| rs12052878 | 2 | 238227594 | G | A | 0.69 | 1.00 | -0.051 | 4.30E-10 | COL6A3 |  |
| rs11712199 | 3 | 14858226 | G | A | 0.91 | 0.99 | 0.081 | 6.80E-12 | FGD5 |  |
| rs370408735 | 3 | 41870038 | C | T | 0.87 | 0.90 | -0.100 | 2.10E-18 | ULK4 | Guo *et al* 2016 |
| rs545996255 | 3 | 58100423 | G | GT | 0.70 | 0.97 | 0.061 | 8.30E-15 | FLNB |  |
| rs2306272 | 3 | 66434643 | T | C | 0.71 | 1.00 | -0.046 | 2.60E-08 | LRIG1 |  |
| rs55914222 | 3 | 128202943 | G | C | 0.97 | 0.99 | 0.190 | 1.40E-16 | GATA2 |  |
| rs13128814 | 4 | 146801002 | G | A | 0.48 | 0.99 | -0.041 | 3.70E-08 | ZNF827 |  |
| rs67846163 | 4 | 174656889 | A | G | 0.77 | 0.99 | -0.082 | 1.30E-20 | HAND2 |  |
| rs2897603 | 5 | 81723109 | C | T | 0.79 | 1.00 | 0.064 | 8.20E-12 | ATP6AP1L |  |
| rs4077816 | 5 | 95582494 | A | G | 0.63 | 0.99 | 0.108 | 2.50E-46 | PCSK1 |  |
| rs7702622 | 5 | 122548721 | C | T | 0.78 | 0.99 | 0.067 | 6.40E-15 | PRDM6 | Vasan *et al* 2009 |
| rs496236 | 6 | 11641601 | A | G | 0.46 | 1.00 | 0.038 | 1.40E-08 | ADTRP |  |
| rs1630736 | 6 | 12295987 | C | T | 0.54 | 0.99 | -0.054 | 1.70E-12 | EDN1 |  |
| rs146170154 | 6 | 36646768 | C | CTA | 0.80 | 0.98 | -0.056 | 3.00E-10 | CDKN1A |  |
| rs1570350 | 6 | 143592386 | A | G | 0.55 | 0.99 | -0.063 | 2.80E-17 | AIG1 |  |
| rs13203975 | 6 | 152333104 | G | A | 0.89 | 0.99 | 0.079 | 5.30E-12 | ESR1 |  |
| rs79215950 | 7 | 35277067 | G | A | 0.62 | 1.00 | 0.062 | 6.60E-17 | TBX20 |  |
| rs6943980 | 7 | 73424373 | A | C | 0.55 | 1.00 | -0.121 | 3.90E-64 | ELN |  |
| rs1583081 | 7 | 85034227 | G | T | 0.58 | 1.00 | -0.079 | 7.00E-28 | SEMA3D |  |
| rs2921059 | 8 | 8317887 | G | T | 0.56 | 0.98 | -0.041 | 3.20E-08 | SGK223 |  |
| rs10097870 | 8 | 11444516 | G | A | 0.54 | 0.99 | -0.049 | 4.70E-12 | GATA4 |  |
| rs11785562 | 8 | 23391493 | G | A | 0.80 | 0.97 | -0.050 | 9.60E-09 | SLC25A37 |  |
| rs9721183 | 8 | 75781818 | C | T | 0.63 | 0.95 | 0.059 | 8.30E-14 | PI15 |  |
| rs16876090 | 8 | 108363596 | G | A | 0.91 | 0.99 | -0.080 | 1.10E-10 | ANGPT1 |  |
| rs562291939 | 8 | 120709336 | T | C | 1.00 | 0.80 | 0.798 | 2.40E-23 | ENPP2 |  |
|  | 8 | 122634926 | CA | C | 0.67 | 0.93 | 0.057 | 3.10E-12 | HAS2 |  |
| rs34557926 | 8 | 124607159 | C | T | 0.63 | 0.99 | -0.071 | 1.10E-21 | FBXO32 |  |
| rs4978966 | 9 | 113662374 | C | T | 0.79 | 1.00 | 0.052 | 4.60E-09 | LPAR1 |  |
| rs16916931 | 10 | 63813744 | A | T | 0.69 | 0.98 | 0.047 | 4.60E-09 | ARID5B |  |
| rs10761716 | 10 | 64882300 | C | G | 0.56 | 0.99 | 0.049 | 2.00E-11 | NRBF2 |  |
| rs71482305 | 10 | 96119130 | C | T | 0.84 | 1.00 | 0.089 | 4.20E-20 | NOC3L |  |
| rs1340837 | 10 | 97542035 | A | G | 0.59 | 1.00 | 0.039 | 1.10E-08 | ENTPD1 |  |
| rs10885378 | 10 | 114491924 | T | C | 0.70 | 0.99 | -0.048 | 1.30E-09 | VTI1A |  |
| rs77889556 | 11 | 17498057 | G | A | 0.83 | 0.91 | -0.070 | 1.20E-12 | ABCC8 |  |
| rs3741025 | 11 | 30851976 | C | T | 0.43 | 0.99 | 0.042 | 2.80E-08 | DCDC1 |  |
| rs111412755 | 11 | 69819139 | C | T | 0.91 | 0.98 | -0.096 | 6.60E-16 | ANO1 | Wild *et al* 2017 |
| rs4936098 | 11 | 130280667 | A | G | 0.37 | 0.98 | -0.053 | 7.40E-13 | ADAMTS8 |  |
| rs2307024 | 12 | 22005003 | T | G | 0.59 | 0.99 | 0.059 | 1.30E-14 | ABCC9 |  |
|  | 12 | 62817410 | TC | T | 0.90 | 0.99 | -0.081 | 1.90E-11 | USP15 |  |
| rs58899389 | 12 | 94199513 | T | C | 0.65 | 0.99 | 0.041 | 4.80E-08 | CRADD |  |
| rs7994761 | 13 | 22871446 | A | G | 0.78 | 0.99 | 0.120 | 1.80E-41 | FGF9 |  |
| rs733166 | 14 | 94464432 | A | G | 0.47 | 1.00 | 0.059 | 1.60E-15 | ASB2 |  |
| rs16970633 | 15 | 40642877 | G | T | 0.83 | 1.00 | -0.052 | 3.70E-08 | PHGR1 |  |
| rs1848050 | 15 | 48862043 | G | A | 0.90 | 0.99 | -0.078 | 4.40E-10 | FBN1 | LeMaire *et al* 2011, Guo *et al* 2016, van 't Hof *et al* 2016 |
| rs6494904 | 15 | 71609522 | G | A | 0.28 | 1.00 | -0.064 | 8.10E-15 | THSD4 |  |
|  | 16 | 56221642 | AAAT | A | 0.56 | 0.98 | -0.047 | 1.20E-11 | GNAO1 |  |
| rs757590420 | 16 | 66896747 | T | C | 1.00 | 0.64 | 0.587 | 1.70E-10 | NAE1 |  |
| rs181062531 | 16 | 69319941 | C | T | 1.00 | 0.31 | 0.667 | 2.70E-08 | SNTB2 |  |

Pirruccello, *et al* — Genetics of the human aorta
| SNP | CHR | BP | Effect Allele | Other Allele | EAF | INFO | BETA | P Value | Nearest Gene | Prior |
| --- | --- | --- | --- | --- | --- | --- | --- | --- | --- | --- |
| rs62053262 | 16 | 69969299 | C | G | 0.95 | 0.99 | 0.207 | 7.40E-36 | WWP2 |  |
| rs7500448 | 16 | 83045790 | A | G | 0.75 | 0.98 | -0.050 | 2.60E-09 | CDH13 |  |
| rs16965180 | 16 | 88989862 | A | G | 0.65 | 0.99 | 0.071 | 1.20E-19 | CBFA2T3 |  |
|  | 17 | 2088848 | CCAGA | C | 0.68 | 1.00 | -0.071 | 1.70E-20 | SMG6 | Vasan *et al* 2009, Wild *et al* 2017 |
| rs4569330 | 17 | 12180624 | G | A | 0.27 | 0.97 | 0.087 | 5.70E-25 | MAP2K4 |  |
|  | 17 | 16155380 | CTTT | C | 0.55 | 1.00 | 0.041 | 4.60E-08 | PIGL |  |
| rs6505216 | 17 | 29206421 | G | T | 0.77 | 0.92 | 0.062 | 8.70E-12 | ATAD5 |  |
| rs8091434 | 18 | 46312960 | C | G | 0.87 | 1.00 | 0.061 | 1.50E-08 | CTIF |  |
| rs3063286 | 20 | 10488552 | T | TTA | 0.47 | 0.94 | 0.050 | 5.60E-12 | SLX4IP |  |
| rs17812022 | 20 | 19007099 | C | T | 0.91 | 1.00 | -0.072 | 7.20E-09 | SLC24A3 |  |
| rs4402860 | 22 | 40554445 | A | T | 0.80 | 1.00 | 0.065 | 1.40E-13 | TNRC6B |  |
| rs755131301 | X | 10401828 | G | A | 0.99 | 0.88 | 0.160 | 3.00E-08 | MID1 |  |

B: Descending thoracic aorta
| SNP | CHR | BP | Effect Allele | Other Allele | EAF | INFO | BETA | P Value | Nearest Gene | Prior |
| --- | --- | --- | --- | --- | --- | --- | --- | --- | --- | --- |
| rs527725 | 1 | 201752429 | A | C | 0.60 | 0.97 | 0.049 | 5.20E-11 | NAV1 |  |
| rs7255 | 2 | 20878820 | T | C | 0.45 | 1.00 | 0.055 | 1.50E-13 | GDF7 |  |
| rs181737440 | 2 | 75328928 | A | G | 1.00 | 0.61 | -0.547 | 1.70E-08 | TACR1 |  |
| rs7580831 | 2 | 238219499 | C | A | 0.68 | 1.00 | -0.048 | 2.30E-09 | COL6A3 |  |
| rs6780370 | 3 | 58074846 | G | A | 0.69 | 0.98 | 0.048 | 5.40E-09 | FLNB |  |
| rs698099 | 3 | 186987941 | G | A | 0.17 | 1.00 | 0.077 | 1.40E-13 | MASP1 |  |
| rs60991988 | 4 | 54801228 | T | G | 0.89 | 0.99 | -0.066 | 2.70E-08 | FIP1L1 |  |
| rs6532500 | 4 | 95580645 | A | G | 0.44 | 0.98 | -0.045 | 1.40E-09 | PDLIM5 |  |
| rs9285863 | 5 | 108071655 | T | C | 0.66 | 0.99 | -0.045 | 5.40E-09 | FER |  |
| rs9263708 | 6 | 31095270 | T | C | 0.74 | 1.00 | -0.054 | 5.90E-11 | PSORS1C1 |  |
| rs77393224 | 6 | 34381136 | G | A | 0.94 | 1.00 | -0.091 | 2.50E-08 | RPS10-NUDT3 |  |
| rs733590 | 6 | 36645203 | T | C | 0.65 | 1.00 | -0.050 | 2.00E-10 | CDKN1A |  |
| rs9362083 | 6 | 85396119 | T | A | 0.42 | 1.00 | 0.046 | 7.80E-10 | TBX18 |  |
| rs4707174 | 6 | 85987918 | A | C | 0.70 | 0.98 | -0.045 | 3.30E-08 | NT5E |  |
| rs2107595 | 7 | 19049388 | G | A | 0.85 | 0.99 | 0.100 | 2.90E-23 | TWIST1 |  |
| rs17774023 | 8 | 10626333 | T | C | 0.69 | 1.00 | 0.048 | 4.50E-09 | PINX1 |  |
| rs532252660 | 8 | 120587297 | C | T | 1.00 | 0.79 | 0.480 | 2.60E-08 | ENPP2 |  |
| rs10740811 | 10 | 30167754 | G | A | 0.41 | 1.00 | 0.083 | 1.10E-27 | SVIL |  |
| rs2797983 | 10 | 95899646 | G | C | 0.45 | 1.00 | -0.072 | 2.10E-20 | PLCE1 |  |
| rs112712475 | 11 | 117065772 | C | A | 0.94 | 0.99 | -0.102 | 4.10E-11 | SIDT2 |  |
| rs4759275 | 12 | 57525756 | G | A | 0.58 | 1.00 | 0.049 | 2.10E-11 | STAT6 | Guo *et al* 2016 |
| rs10744777 | 12 | 112233018 | T | C | 0.66 | 1.00 | -0.048 | 1.60E-09 | ALDH2 |  |
| rs12885183 | 14 | 21545230 | A | G | 0.77 | 0.99 | 0.054 | 7.70E-10 | ARHGEF40 |  |
| rs7143356 | 14 | 23881083 | T | C | 0.62 | 1.00 | 0.045 | 1.80E-08 | MYH6 |  |
| rs12890024 | 14 | 94469801 | A | G | 0.62 | 0.98 | 0.054 | 1.90E-12 | OTUB2 |  |
| rs71472433 | 15 | 40649609 | A | C | 0.83 | 0.99 | -0.063 | 1.30E-09 | DISP2 |  |
| rs17352842 | 15 | 48694211 | C | T | 0.81 | 1.00 | -0.060 | 2.10E-10 | FBN1 | LeMaire *et al* 2011, Guo *et al* 2016, van 't Hof *et al* 2016 |
| rs1048661 | 15 | 74219546 | G | T | 0.66 | 0.99 | -0.050 | 1.70E-10 | LOXL1 | Vasan *et al* 2009 |
| rs8182076 | 15 | 79072988 | C | T | 0.58 | 0.99 | -0.043 | 1.20E-08 | ADAMTS7 |  |
| rs62053262 | 16 | 69969299 | C | G | 0.95 | 0.99 | 0.119 | 3.30E-12 | WWP2 |  |
| rs56014161 | 17 | 77910740 | C | T | 0.69 | 0.99 | -0.044 | 2.70E-08 | TBC1D16 |  |
| rs12607403 | 18 | 46343221 | C | T | 0.12 | 0.99 | -0.078 | 3.40E-12 | CTIF |  |
| rs8102624 | 19 | 2161443 | G | A | 0.92 | 1.00 | 0.101 | 3.30E-13 | DOT1L |  |
| rs2303040 | 19 | 39138608 | T | C | 0.51 | 0.99 | -0.049 | 2.20E-10 | ACTN4 |  |
|  | X | 114835773 | TA | T | 0.43 | 0.92 | -0.036 | 8.60E-09 | PLS3 |  |

imputation INFO score. BETA = effect size per effect allele on the inverse normal transformed trait. Prior = known from prior publications addressing common genetic variation linked to aortic size, aortic aneurysm, or aortic dissection<sup>15–19</sup>.

**Figure 2:**
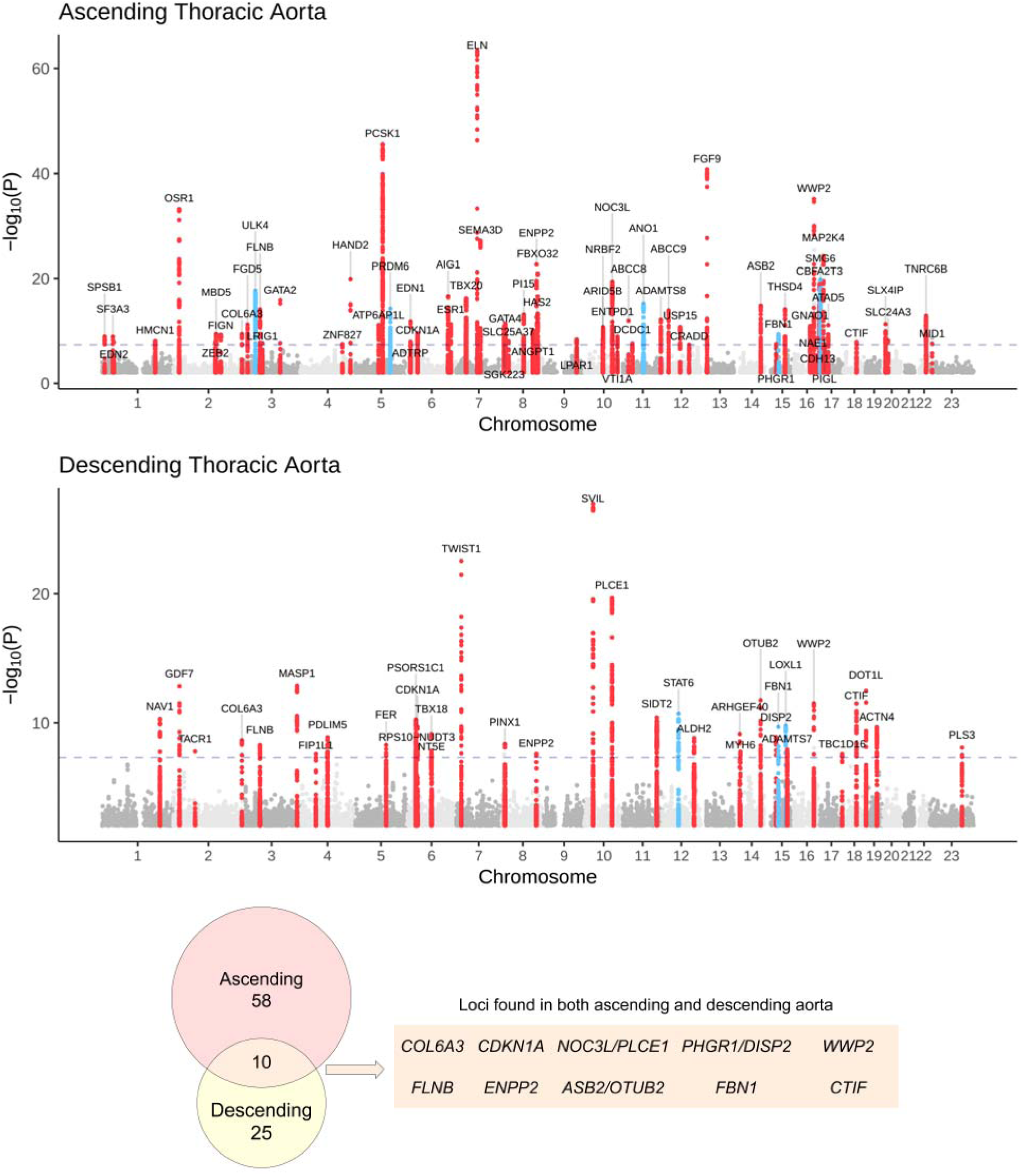
Genome-wide association study results. **Panels A & B**: Loci with P < 5×10^−8^ are shown in red (if not previously reported) or blue (if previously reported in common variant association studies for aortic size or disease status [aneurysm or dissection]). The X chromosome is represented as ‘23’. **Panel C**: Venn diagram showing the number of loci uniquely associated at P < 5×10^−8^ with either the ascending or descending thoracic aorta. Those in orange are associated with both and are enumerated in the table to the right. Loci whose lead SNP’s nearest gene differs between ascending and descending are demarcated as “Ascending/Descending”.

Previous analyses of thoracic aortic phenotypes including aortic root diameter, ascending aortic dissection, or thoracic aortic aneurysm have identified only 15 genome-wide significant loci to date; of these, seven achieved genome-wide significance in our study, including all three loci that have been associated with thoracic aortic dissection (near *FBN1, ULK4*, and the *STAT6*/*LRP1* locus; **Supplementary Table 5**)^15–19^.

We sought to replicate our GWAS findings in 3,287 participants from the Framingham Heart Study (FHS) who had genotyping data and cross-sectional imaging of the ascending and descending thoracic aorta by computed tomography^20,21^. Since the FHS sample size was an order of magnitude smaller than our discovery population in the UK Biobank, we focused on directional agreement. Of the 67 autosomal lead SNPs in the ascending aorta, 54 were identified in the FHS dataset. 44 of these 54 SNPs were directionally consistent in both datasets (two-tailed binomial P = 3.4×10^−6^; **Supplementary Figure 4A**). 30 of the 34 autosomal lead SNPs from the descending aorta were identified in FHS, and 27/30 were directionally consistent (binomial P = 8.4×10^−6^; **Supplementary Figure 4B, Supplementary Table 6**). Thus, despite comprising a significantly smaller sample, as well as using a different imaging modality and measurement technique, the FHS results were aligned with our findings from the UK Biobank.

We used genetic correlation to gain insight into the relationship between aortic diameter and other cardiovascular and anthropometric phenotypes. In the UK Biobank, the ascending and descending aortic phenotypes had a genetic correlation with one another of 0.47 (95% CI 0.43-0.51) as estimated by BOLT-REML^22,23^. We used linkage disequilibrium (LD) score regression to assess genetic correlation between the aortic traits and 272 quantitative phenotypes from the UK Biobank that were precomputed by the Neale Lab^24,25^, linking aortic size to measures of height, weight, and blood pressure, among other traits. As expected, we observed positive genetic correlations between aortic size and anthropometric measures such as height and weight, as well as related phenotypes such as blood pressure (**Supplementary Table 7**; **Supplementary Figures 5-6**).

To gain more insight into the GWAS loci themselves, we then took three approaches to prioritize genes at each locus and to link those genes to relevant cell types. First, we conducted a transcriptome wide association study (TWAS), linking predicted gene expression in aorta (based on GTEx v7) with aortic size (**Figure 3A**)^26,27^. We identified a total of 51 genes that were significantly associated with the dimensions of the ascending or descending aorta at P < 5×10^−8^. The strongest TWAS associations in the ascending aorta included *ULK4*, a gene previously linked with aortic dissection, and *THSD4*, whose protein product binds to fibrillin (*FBN1*) and modulates microfibril assembly^28^. In addition to *THSD4* and *FBN1*, several other GWAS loci harbored genes involved in the process of elastogenesis including *LOXL1* and the gene encoding elastin itself, *ELN*. The strongest TWAS association in the descending aorta was with the gene *SVIL*, in which increased transcription was associated with increased aortic diameter (**Figure 3A**).

**Figure 3:**
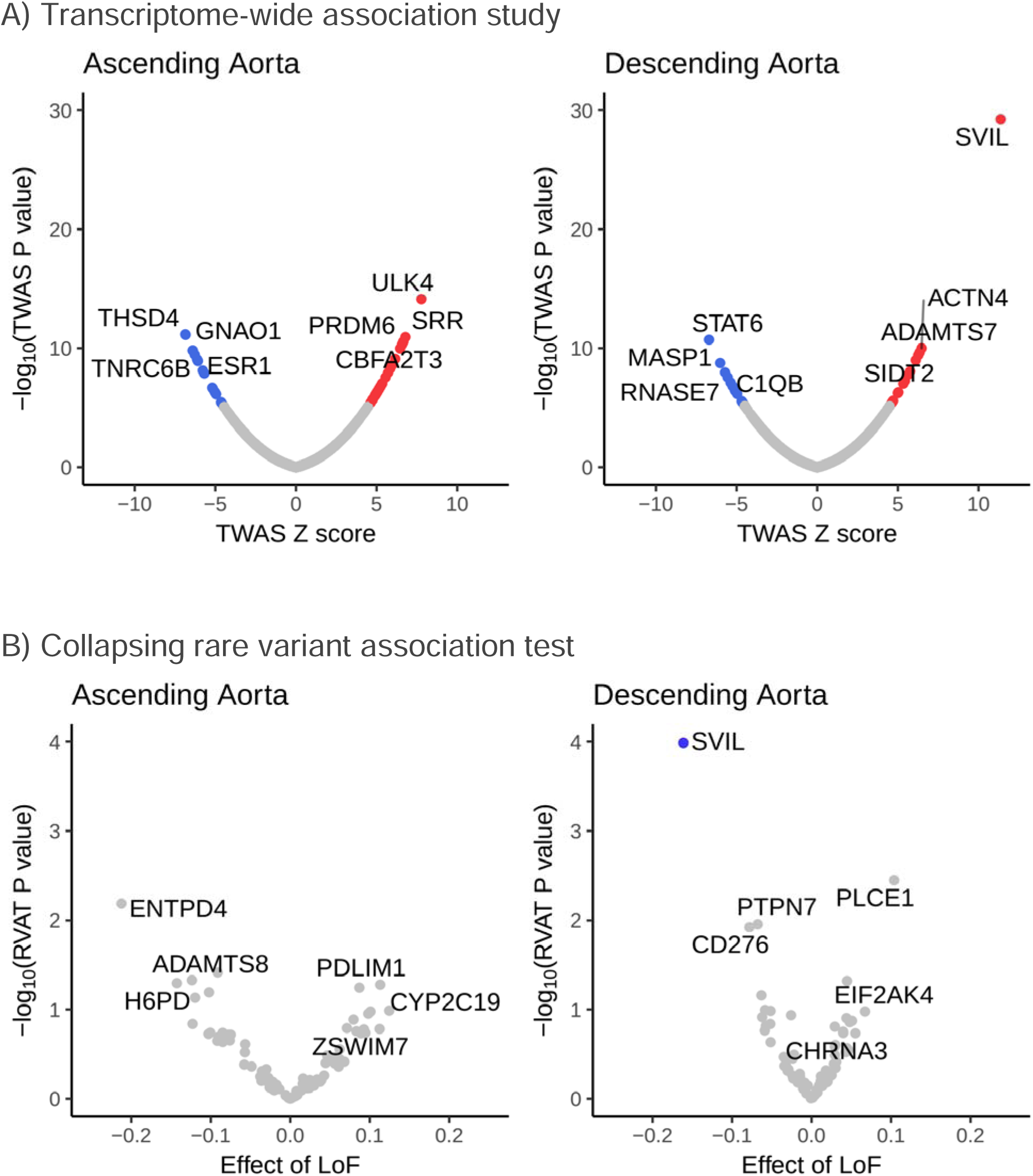
Gene-level association tests. **Panel A**: Protein-coding genes associated with the size of the ascending (**left panel**) and descending (**right panel**) thoracic aorta based on an integrated gene expression prediction are shown. The **x axis** represents the magnitude of the TWAS Z score, while the **y axis** represents the −log10 of the TWAS P value. Traits achieving Bonferroni significance are colored red (positive correlation) or blue (negative correlation). The top 4 positively and negatively correlated traits are labeled. **Panel B**: Rare variant collapsing burden test results are depicted. Loss of function carrier status in each gene was tested for association with the size of the ascending (**left panel**) and descending (**right panel**) thoracic aorta. The **x axis** represents the effect size of LoF in each gene on aortic size, while the **y axis** represents the −log10 of the association P value in a logistic model. The one gene achieving Bonferroni significance (*SVIL*) is colored blue for its negative correlation with the size of the descending thoracic aorta. The top 3 genes are labeled.

Second, we conducted a rare variant association test in over 12,000 UK Biobank participants with both aortic imaging and exome sequencing data (**Figure 3B**). We found that rare, loss-of-function variation in one gene, *SVIL*, was significantly associated with a reduced diameter of the descending aorta (14 carriers; loss-of-function effect size −0.16cm, 95% CI −0.08 to −0.24 cm, P=1.03×10^−4^).

Third, we undertook direct analysis of tissue and cell-specific expression patterns to localize and identify relevant cell types. We used tissue-specific LD score regression to test for enrichment of the aortic diameter GWAS results in 53 GTEx v6 tissue types^27,29^. As expected, for the ascending aortic loci, enrichment was confirmed in aortic and coronary artery tissues (P=1.5×10^−4^ and P=4.7×10^−4^, respectively); for the descending aorta, enrichment was confirmed in aortic tissue only (P=6.4×10^−4^; **Supplementary Tables 10-11**). These data are consistent with the expectation that the aorta itself is the most relevant tissue linked with our findings. Therefore, we incorporated an analysis of single-nucleus RNA sequencing (snRNA-seq) of rat aorta to identify potentially relevant cell types for the genes at aortic GWAS loci. We sequenced the transcriptomes of 18,707 single nuclei and identified 10 primary cell clusters in the rat aorta (**Figure 4A**). Through comparison of unique transcriptional profiles in each cluster to canonical cell markers, we identified populations comprising vascular smooth muscle cells, fibroblasts, three distinct types of endothelial cells and two types of adipocytes (**Figure 4B**). We then examined the cell type-specific expression of the genes prioritized by the TWAS (**Figure 4C and 4D**).

**Figure 4:**
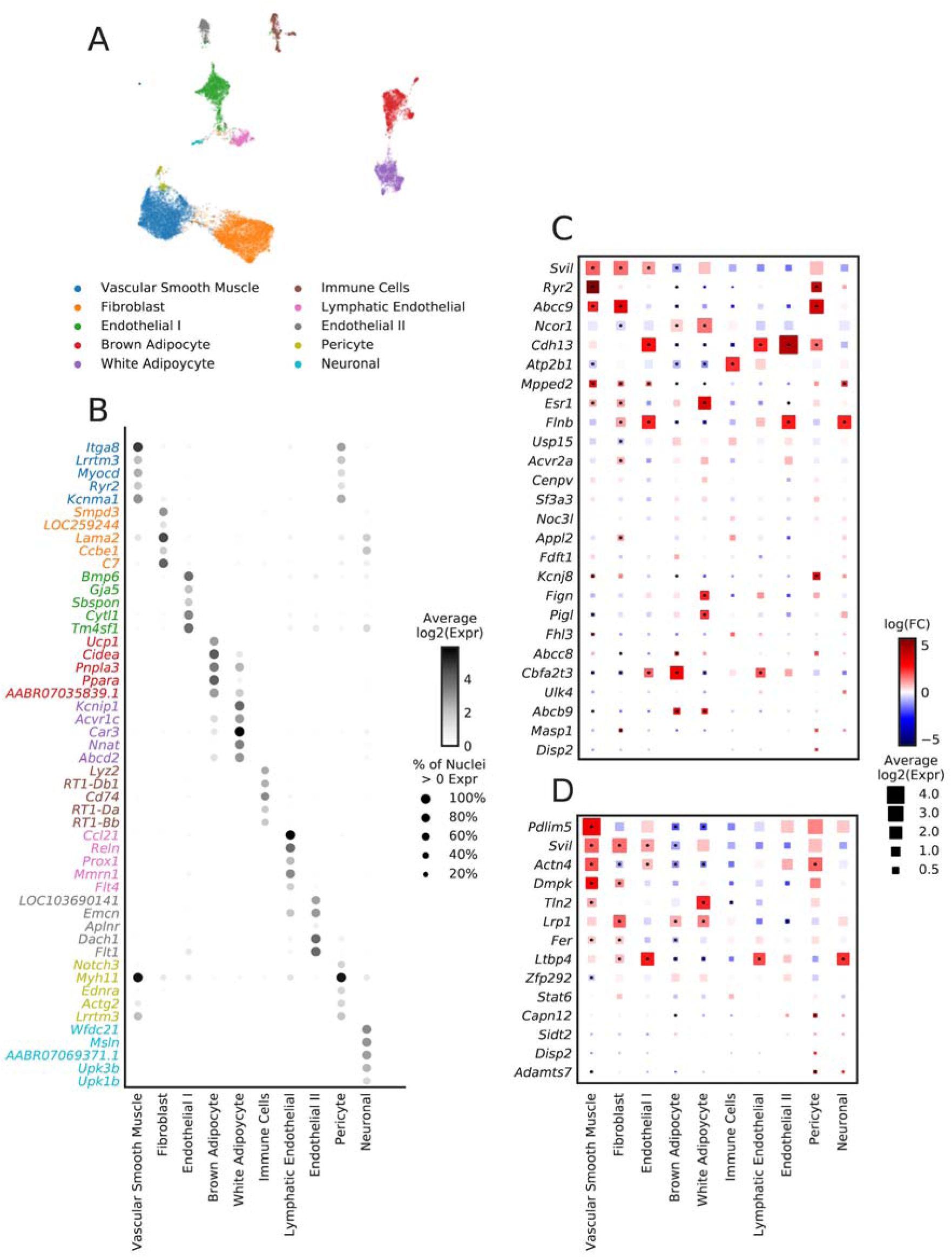
Single nucleus RNA sequencing analyses in rat aorta. Single nucleus RNA-seq was performed on aorta tissue from 6 Wistar rats. **Panel A**: Uniform manifold approximation and projection (UMAP) revealed 10 main clusters. Each dot represents an individual nucleus, colored and labeled by putative cell type as identified from Louvain clustering. **Panel B**: The top five most selectively expressed genes for each cluster were identified as those with the largest fold-change difference in expression comparing the given cluster with all other clusters, only considering genes expressed in at least 10% of nuclei and with a Benjamini-Hochberg corrected p < 0.01. The shade of the dot represents the average log2 expression for a gene across all nuclei in a given cluster and the size of the dot represents the percentage of nuclei in the cluster with non-zero expression. The cell type labels were created by comparing selectively expressed genes in each cluster of nuclei with the literature. **Panels C and D**: Cell-type specificity of genes with expression data supported by the TWAS in the ascending (**Panel C**) and descending (**Panel D**) aorta. The size of each square represents the average log2(Expr) for a gene across all nuclei in a given cluster. The color represents the log fold-change comparing the expression of the given gene in each cluster to all other clusters based on a formal differential expression model. A dot represents significant up- or down-regulation in the given cluster based on a Benjamini-Hochberg correction for multiple testing at FDR < 0.01. Expr = Normalized nucleus-level expression calculated as the number of counts of a gene divided by the total number of counts in the nucleus and multiplied by 10,000; FC = Fold-change.

Remarkably, a SNP near the *SVIL* locus was the strongest GWAS signal for the descending aorta, and *SVIL* was the gene most strongly associated in the TWAS (increased expression is linked to a larger descending aorta diameter; **Figure 3A, Supplementary Tables 12-13**), as well as the strongest association signal in the rare variant association test in which loss of function is linked to a smaller descending aorta (**Figure 3B, Supplementary Table 14**). snRNA-seq revealed that *SVIL* is most strongly expressed in vascular smooth muscle cells within the aorta(**Figure 4C and 4D**), consistent with a role in aortic size determination. *SVIL* encodes the protein supervillin, an F-actin and myosin II binding protein that localizes to and coordinates the action of cell surface extensions called ‘invadosomes’. These promote matrix degradation through the localized release of extracellular matrix-lytic enzymes such as disintegrin- and-metalloprotease domain-containing proteins and matrix metalloproteinases^30,31^.

Our genetic and single-nucleus transcriptomic analyses also highlight *WWP2*, which is linked to the size of both ascending and descending aorta. The lead SNP (rs62053262) is an expression quantitative trait locus (eQTL) in the aorta for *WWP2*^27^; the rs62053262 G allele corresponds to reduced expression of WWP2 in aorta and smaller aortic size. *WWP2* acts as an E3 ubiquitin ligase for PTEN^32^ and has previously been shown to regulate cardiac fibrosis through modulation of SMAD signaling^33^. Examining rat single-nucleus expression data, we show that *WWP2* expression is enriched in aortic vascular smooth muscle cells (**Supplementary Figure 7**).

In other cardiovascular phenotypes, GWAS loci have been enriched for Mendelian genes^34,35^, so we asked whether the loci identified in our study were in closer proximity to more genes implicated in Mendelian aortopathies than expected by chance. We did not find an enrichment of previously described Mendelian thoracic aortic aneurysm and dissection genes^36^ (23 genes; 2 overlapping with ascending loci, P=0.09; 1 overlapping with descending loci, P=0.27 by one-tailed permutation tests). However, our analysis has independently identified loci containing relevant genes such as *FBN1*, well described as the causal gene in Marfan syndrome^37^, and loci near genes such as *PI15*, known to cause arterial dysfunction in rats^38^, and *ABCC9*, a rare recessive cause of aortic aneurysm in humans^39^. Other loci suggest the involvement of novel genes within networks previously implicated in aortic disease; for instance, the protein product of *ASB2* is part of the E3 ligase that targets both filamin B (encoded by *FLNB*, the nearest gene to a lead SNP on chromosome 3) and the known aortic disease protein filamin A (*FLNA*) for degradation^40^. Moreover, TGF-β signaling, heavily implicated in clinical aortic disease, is also represented in our GWAS gene set as indicated by MAGMA analysis (**Supplementary Figure 8**; **Supplementary Tables 8-9**)^41^.

Finally, we probed the clinical relevance of the GWAS loci by asking whether a polygenic score for ascending aortic size produced from these loci was associated with aortic disease risk. We analyzed the remaining UK Biobank participants who had not undergone MRI and who did not have a diagnosis of aortic disease at enrollment. A polygenic score from the 83 autosomal, independently significant SNPs from the ascending aorta GWAS was strongly associated with the 381 incident cases of aortic aneurysm or dissection (HR = 1.44 per standard deviation; CI 1.30-1.59; P = 3.7×10^−12^). Participants in the top 10% of the polygenic score had a 2.2-fold hazard ratio compared to the remaining 90% of the cohort (CI 1.7-2.9; P = 5.2×10^−10^; **Figure 5**).

**Figure 5:**
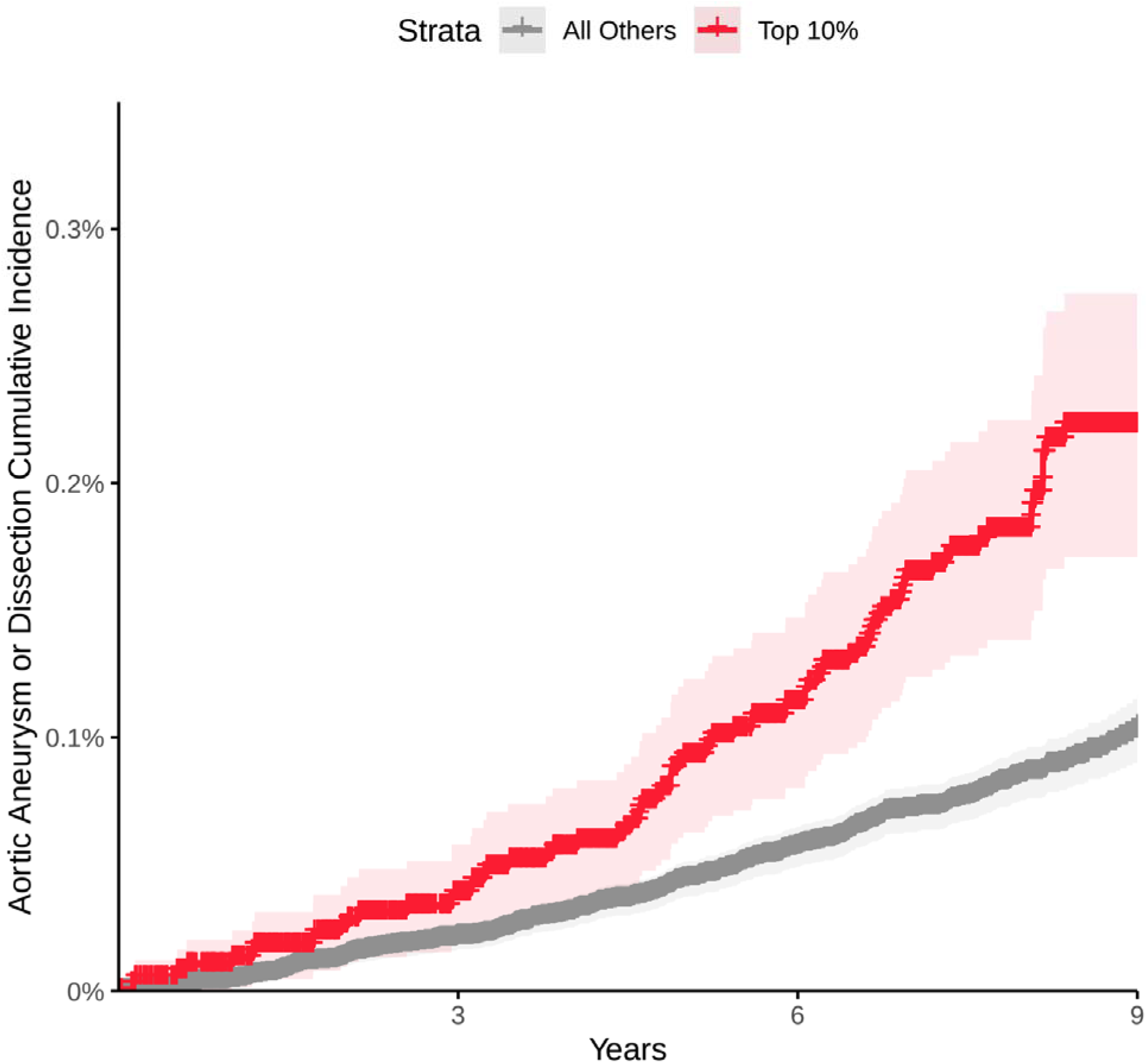
Cumulative incidence of thoracic aortic aneurysm or dissection stratified by polygenic score. The cumulative incidence (1 minus the Kaplan-Meier survival estimate) of a diagnosis of aortic aneurysm or dissection (Y axis) is plotted against the number of years since UK Biobank enrollment (X axis). Individuals in the top tenth percentile of the polygenic score for ascending aorta size are shown in red; the remaining 90% are shown in gray. The 95% confidence intervals (from the cumulative hazard standard error) are represented with lighter colors.

Our study is subject to several limitations. The study population largely consisted of European-ancestry UK Biobank participants, limiting generalizability to other populations. The aortic measurements were derived from a deep learning model that was trained on cardiologist-annotated segmentation data, but the vast majority of images were not manually reviewed; nevertheless, genetic results derived from manually annotated FHS imaging data were generally concordant with our findings. Whereas genetic conservation between the rat and human is high, single-nucleus RNA expression data from the rat, as for other model organisms, are imperfect representations of the human aorta. Finally, because thoracic aortic aneurysm is not routinely assessed in screening tests, the effect estimate of the ascending aortic polygenic score is likely to be biased due to ascertainment in UK Biobank participants; future analyses in external datasets will be required to confirm the observation linking the polygenic score to incident aortic aneurysm or dissection.

In summary, we used deep learning to assess the size of the ascending and descending thoracic aorta using magnetic resonance imaging data in a large population-based biobank. We identified 63 novel loci in the ascending aorta and 32 in the descending aorta, explored their relationships to other traits, and assessed their association with aortic aneurysm or dissection. These findings permit several conclusions. First, these results demonstrate that deep learning is a powerful tool for deriving quantitative phenotypes from raw signal data at a population level. In particular, by using transfer learning from a deep learning model trained on a large but unrelated set of images compiled for a different task, we were able to develop a useful model while manually annotating only a small number of images. Second, these results highlight the value of studying quantitative traits, such as aortic size, in order to gain greater understanding of disease processes underlying aneurysm and dissection. Third, the modest genetic correlation and limited locus overlap of the ascending and descending thoracic aorta highlight their distinct biology. Fourth, we prioritize several potential gene targets based on integration of GWAS, TWAS, and rare variant analyses, and identify their likely cell type of relevance with snRNA-seq. Fifth, a polygenic score for ascending aortic size is an independent risk factor for aneurysmal enlargement of aorta. In the future, it will be interesting to determine if a model incorporating a polygenic score and clinical risk factors might identify high-risk, asymptomatic individuals who would benefit from thoracic imaging to screen for ascending aortic aneurysm.

## Online Methods

### Study design

The UK Biobank is a richly phenotyped, prospective, population-based cohort that recruited 500,000 individuals aged 40-69 in the UK via mailer from 2006-2010^42^. In total, we analyzed 487,283 participants with genetic data who had not withdrawn consent as of October 2018. Access was provided under application #7089. Analysis was approved by the Partners HealthCare institutional review board (protocol 2013P001840). GWAS replication was performed in an imaging substudy of the community-based Framingham Heart Study (FHS) Offspring and Third-Generation cohorts; participants were ascertained based on sex-specific age cutoffs (≥ 35 years for men and ≥ 40 years for women), and weight < 350 pounds as described previously and approved by the institutional review boards of the Boston University Medical Center and the Massachusetts General Hospital^20^.

A deep learning model for aorta pixel recognition in cardiac MRI was developed and applied to imaging data from UK Biobank participants. Genetic discovery of loci related to ascending and descending thoracic aortic size was performed in this cohort. A replication GWAS was performed in FHS. A transcriptome-wide association study (TWAS) and rare-variant association tests were performed to prioritize genes at each genomic locus, and we analyzed single-nucleus gene expression in rat aortas in order to identify relevant cell types for these genes. A polygenic score produced from the GWAS SNPs was used to predict incident aortic disease diagnosis in the remaining UK Biobank participants who had not undergone cardiac MRI.

### Cardiac magnetic resonance imaging

The UK Biobank is conducting an imaging substudy on 100,000 participants which is currently underway^43,44^. Cardiac magnetic resonance imaging was performed with 1.5 Tesla scanners (MAGNETOM Aera, Siemens Healthcare), using electrocardiographic gating for cardiac synchronization^44^. A balanced steady-state free precession cine, consisting of a series of exactly 100 images throughout the cardiac cycle, was acquired for each participant at the level of the right pulmonary artery^44^. In total, 3,840,140 images from 37,910 UK Biobank participants were analyzed. Of these, 458 participants had one or more repeat sets of images, and four had incomplete studies with fewer than 100 images.

### Deep learning for segmentation of the aorta

Segmentation maps were traced for the ascending and descending thoracic aorta manually by a cardiologist (JPP). To produce the final model used in this manuscript, 116 samples were chosen at random, manually segmented, and were used to train a deep learning model with fastai v1.0.59^45^. The model consisted of a U-Net-derived architecture, where the encoder was a resnet34 model pre-trained on ImageNet^14,45–48^. 80% of the samples were used to train the model, and 20% were used for validation. Development versions prior to this final model are detailed in the subsequent section.

During training, all images were resized to be 120 pixels in width by 98 pixels in height for the first half of training, and then 240 pixels in width by 196 pixels in height for the second half, detailed below. The Adam optimizer was used, and the model was trained with a minibatch size of 8 (when training with half-dimension images) 4 (when training with full-dimension images)^49^. Rather than using extensive hyperparameter tuning, the model was instead trained using a cyclic learning rate training policy, which alternately decreases and increases the learning rate during training^50^. For the first half of training using half-dimension images, the maximum learning rate (the step size during gradient descent) was set at 0.001, with 40% of the iterations permitted to have an increasing learning rate during each epoch across 20 epochs. This was performed while keeping all ImageNet-pretrained layers fixed, so that only the final layer was fine-tuned. Then all layers were unfrozen and the model was trained for an additional 15 epochs with the same maximum learning rate. For the second half of training using full-dimension images, the maximum learning rate was set to 0.0002, with 30% of the iterations permitted to have an increasing learning rate. Then, all layers were unfrozen and the model was trained for an additional 15 epochs with a maximum learning rate of 0.00002.

Throughout training, augmentations (random perturbations of the images) were applied as a regularization technique. These augmentations included affine rotation, zooming, and modification of the brightness and contrast. Because medical imaging data is not symmetric across the midline of the human body, we did not permit mirroring transformations. 92 images were used to train the model, and 24 were held out for validation; the model achieved 97.4% pixel categorization accuracy in the held-out validation set.

This model was then used to infer segmentation of the ascending and descending aorta on all 3,840,140 images in the dataset. During inference, adaptive pooling was used to permit arbitrary image sizes^51^, which allowed us to produce output that matched the input size and thereby preserve the number of millimeters per pixel as reported in the DICOM metadata.

### Development versions of the deep learning model

The first batch of manual segmentation mapping of aorta was performed by one cardiologist (JPP) on 58 images, a sample size that was chosen to balance the time required for annotation (approximately 5 minutes per sample) against the need for diverse data to train the ImageNet-based segmentation model. A deep learning model (with the same training parameters as described above) trained with this data (using 47 images for training and 11 for validation) achieved 95.1% pixel accuracy.

When the output of this model was visualized, the notable recurring error was the miscategorization of breast implants as aorta. To produce the final training set, the sample size was doubled from 58 images to 116, of which 15 had breast implants. No other significant hyperparameter tuning was performed.

### Segmentation quality control

As the cine videos consisted of 100 still frame images, single-image quality control was performed first. Images which lacked any pixels labeled as aorta were excluded. Next, the connected components labeled as ascending or descending aorta were counted using the Rosenfeld-Pfaltz algorithm^52^. Images having a number of aortic components more than 5 standard deviations above the mean were excluded. Any participant with at least one image that failed this quality control procedure was excluded from further analysis.

Then, we performed a quality control step that took advantage of the dynamics of the cardiac cycle. We computed the largest frame-to-frame change in the cross-sectional area of the ascending and descending aorta. Outliers beyond 10 standard deviations above the mean were excluded. Then, samples were excluded if the variance in the number of components of the ascending or descending aorta across all frames throughout the cardiac cycle was above 10 standard deviations beyond the mean amount of variance in the full cohort. At the completion of quality control, 34,764 individuals remained for further analysis.

### Extraction of aortic traits

Having identified which pixels represented aorta, we were able to determine the aorta’s cross-sectional dimensions. The aorta was treated as an ellipse: major and minor axes were computed using classical image moment algorithms^53^. For both the ascending and the descending thoracic aorta, the length of the minor elliptical axis (in centimeters) was ascertained at the point in the cardiac cycle when the aorta was at its maximum size (closely corresponding with end-systole). The minor axis was chosen for analysis because imperfection in the orientation of the plane of image acquisition may falsely elongate the apparent major axis of the ascending and descending aorta; in contrast, the dimension of the minor axis is not affected by such perturbations. The length of the minor axis (i.e., the diameter) of the ascending and descending aorta were treated as our primary phenotypes for subsequent analyses.

### Aortic disease codes

International Classification of Diseases version 10 (ICD-10) codes and Office of Population Censuses and Surveys Classification of Interventions and Procedures version 4 (OPCS-4) codes used to define aortic procedures and thoracic aortic aneurysm, dissection, or rupture are detailed in **Supplementary Table 15**. These definitions were, respectively, used for GWAS participant exclusion and polygenic score assessment.

### Correlation between phenotypes and aortic measurements

Statistical analyses were conducted with R version 3.6 (R Foundation for Statistical Computing, Vienna, Austria). We conducted phenome-wide association studies (PheWAS) to assess the relationship between the observed aortic traits and (a) other continuous traits measured in the UK Biobank, and (b) other disease phenotypes based on ICD-10 and OPCS-4 codes.

All 34,764 participants with aortic measurements were used in the continuous trait PheWAS. The number of participants modeled for each trait varied based on availability in the UK Biobank. 674 traits were analyzed using a linear model accounting for the MRI serial number, sex, PC1-5, age at enrollment, the cubic natural spline of age at the time of MRI, and the genotyping array.

The same covariates were used in a logistic regression model testing the relationship between the aortic traits and 1,446 PheCode-defined diseases derived from hospital billing codes. (Because most cardiac MRIs in the UK Biobank were performed near the end of currently available follow-up time, assessment of incident disease after ascertainment of aortic size was not feasible.)

### Genotyping, imputation, and genetic quality control

As detailed previously, UK Biobank samples were genotyped on either the UK BiLEVE or UK Biobank Axiom arrays, then centrally imputed into the Haplotype Reference Consortium panel and the UK10K+1000 Genomes panel^54^. Variant positions were identified using the GRCh37 human genome reference. Genotyped variants with genotyping call rate < 0.95 and imputed variants with INFO score < 0.3 or minor allele frequency <= 0.001 in the analyzed samples were excluded. After variant-level quality control, 16,001,524 imputed autosomal variants and 562,369 imputed variants on the X chromosome remained for analysis.

Participants without imputed genetic data, or with a genotyping call rate < 0.98, mismatch between self-reported sex and sex chromosome count, sex chromosome aneuploidy, excessive third-degree relatives, or outliers for heterozygosity as defined centrally by the UK Biobank were excluded^54^. In addition, we excluded participants with a prior history of aortic repair or other aortic procedures.

### Heritability and genetic correlation of aortic traits

BOLT-REML v2.3.4 was used to assess the SNP heritability of the minor axis length of the ascending and descending thoracic aorta and their genetic correlation with one another using the directly genotyped variants in the UK Biobank^22^.

### Genome-wide association study of aortic traits

We analyzed the minor axis length of the ascending and descending thoracic aorta at the frame within the cardiac cycle when they were at their largest. These traits were first residualized on age at enrollment, the natural spline of age at the time of MRI with 3 knots, the first five principal components of ancestry, sex, the genotyping array, and the MRI scanner’s unique identifier. The residuals were found to be non-normally distributed (with non-zero skewness and kurtosis). Therefore, these residuals were inverse-normal transformed prior to genetic analysis^55^.

Genome-wide association studies for the minor axis length of the ascending and descending thoracic aorta were conducted using BOLT-LMM version 2.3.4 to account for cryptic population structure and sample relatedness^22,23^. We used the full autosomal panel of 713,628 directly genotyped SNPs that passed quality control to construct the genetic relationship matrix (GRM). GWAS covariates included age at enrollment, age and age^2^ at the time of MRI, the first five principal components of ancestry, sex, the genotyping array, and the MRI scanner’s unique identifier. Associations on the X chromosome were also analyzed, using all autosomal SNPs and X chromosomal SNPs to construct the GRM (N=731,238 SNPs), with the same covariate adjustments and significance threshold as in the autosomal analysis. In this analysis mode, BOLT treats individuals with one X chromosome as having an allelic dosage of 0/2 and those with two X chromosomes as having an allelic dosage of 0/1/2. Variants with association P < 5×10^−8^ were considered to be genome-wide significant.

In order to identify independently significantly associated variants, linkage disequilibrium (LD) clumping was performed with plink-1.9^56^ in the same participants used to conduct the GWAS. We used a wide 5-megabase window (--clump-kb 5000) and a stringent LD threshold (--r2 0.01) in order to identify independently significant SNPs despite long LD blocks (particularly on chromosome 16 near *WWP2*). Using the independently significant SNPs, distinct genomic loci were defined by starting with the SNP with the strongest P value, excluding other SNPs within 500kb, and iterating until no SNPs remained. The independently significant SNPs that defined each genomic locus are termed the lead SNPs. Lead SNPs were tested for deviation from Hardy-Weinberg equilibrium at a threshold of P < 1×10^−656^.

### GWAS Replication

The genetic profiles of FHS participants were measured by the Affymetrix GeneChip 500k Array Set & 50K Human Gene Focused Panel, and genotyping was called using BRLMM as previously described^57,58^. Variants with call rate < 0.97, HWE P < 10^−6^, N > 100 Mendelian errors, or MAF < 0.01 were excluded. The remaining variants were then imputed to the TOPMed imputation panel using Michigan Imputation Server (https://imputationserver.sph.umich.edu/index.html)^59^. A multi-detector computed tomography (CT) scanner (General Electric Lightspeed + 8 detector scanner) was used to assess the aorta in FHS participants^20,21^. All measurements have been deposited into dbGaP (Accession: phs000007.v30.p11). The association between each genetic variant and CT traits was tested with linear mixed effects models using the *kinship* package in R, and adjusted for sex, age, age square, cohort, and first five principal components of ancestry.

We then identified lead SNPs from the main GWAS which were also available in the FHS GWAS and ensured that their effect directions were aligned based on effect allele and non-effect allele. We performed a two-tailed binomial test for directional consistency of effect direction, using the null hypothesis that, for each of these independent SNPs, directional agreement would be expected by chance 50% of the time. We then performed linear regression, predicting the FHS Z scores with the UK Biobank Z scores. To assess whether more extreme Z scores corresponded with better agreement between the primary study and the replication study, we modified the SNPs in the linear model by adjusting the UK Biobank SNP P value inclusion threshold from P < 5×10^−6^ to P < 5×10^−14^, and assessed the coefficient of determination of the model at several incremental thresholds within that range. This analysis was performed for both ascending and descending aorta.

### LD score regression for inflation

Linkage disequilibrium (LD) score regression analysis was performed with *ldsc* version 1.0.0^60^. For each GWAS, the genomic control factor (lambda GC) was partitioned into polygenic and inflation components using the software’s defaults.

### Genetic correlation with other quantitative traits

Genetic correlation across traits was assessed using *ldsc*^25^ in 272 continuous traits from the UK Biobank whose *ldsc*-formatted summary statistics were made available by the Neale Lab^24^.

We then applied *aberrant*, a software package in R^61^, to cluster the 272 traits based on their genetic correlation Z scores. Using lambda (the ratio of standard deviations of outliers vs inliers) set to 40, we identified a large inlier cluster and two outlier clusters based on differential genetic correlation with ascending or descending aorta.

### Tissue-specific LD score regression

To address which tissues were most tightly linked to the ascending and descending aorta GWAS results, we applied tissue specific LD score regression against 53 GTEx v6 tissue types that were preprocessed by the *ldsc* authors^27,29^. The *ldsc* authors identified genes that were specifically expressed in each tissue, retaining the top 10% of genes most specifically expressed from each of the 53 tissues. We then conducted stratified LD score regression with these specifically enriched gene sets (*ldsc-SEG*) to determine the contribution of the tissue-specific expression to the heritability of the size of the aorta. The returned P value represents the probability of seeing such a large coefficient if the null hypothesis (that the tissue is not enriched) were true; i.e., it tests whether the tissue-specific contribution is distinguishable from zero. Significance was determined using a false discovery rate (FDR) of 5%.

### Mendelian aortopathy gene set enrichment

We considered the 23 thoracic aortic aneurysm and dissection-related genes from Category A, B, or C from Renard, *et al*, to be Mendelian aortopathy genes^36^. SNPsnap was used to generate 10,000 sets of SNPs that match the lead SNPs from the GWAS based on minor allele frequency, number of SNPs in linkage disequilibrium, distance to the nearest gene, and gene density at the locus ^62^. A lead SNP was considered to be near a Mendelian locus if it was within 500 kilobases upstream or downstream of any gene on the panel. Significance was assessed by permutation testing across the 10,000 SNP sets to determine the neutral expectation for the number of overlapping genes in loci with characteristics similar to ours, yielding a one-tailed permutation P value.

### Transcriptome-wide association study

For both ascending and descending thoracic aorta, we performed a TWAS to identify genes whose imputed cis-regulated gene expression corrrelates with aortic size^26,63–65^. We used *FUSION* with eQTL data from GTEx v7. Precomputed transcript expression reference weights for the aorta (N=6,462 genes) were obtained from the *FUSION* authors’ website (http://gusevlab.org/projects/fusion/)^26,27^. *FUSION* was then run with its default settings.

### MAGMA gene set analysis

We tested 10,992 gene sets from MSigDB for enrichment in the ascending and descending aortic GWAS results using MAGMA 1.07b^41,66^. We used gene locations for GRCh37 and European reference data that were preprocessed by MAGMA’s authors (https://ctg.cncr.nl/software/magma). We used the composite “GO_PANTHER_INGENUITY_KEGG_REACTOME_BIOCARTA” gene sets from MSigDB provided by the MAGENTA authors^67,68^.

### Exome sequencing in UK Biobank

Samples from the UK Biobank were chosen for exome sequencing based on enrichment for MRI data and linked health records^69^. Exome sequencing was performed by Regeneron and reprocessed centrally by the UK Biobank following the Functional Equivalent pipeline^70^. Exomes were captured with the IDT xGen Exome Research Panel v1.0, and sequencing was performed with 75-base paired-end reads on the Illumina NovaSeq 6000 platform using S2 flowcells. Alignment to GRCh38 was performed centrally with BWA-mem. Variant calling was performed centrally with GATK 3.0^71^. Variants were hard-filtered if the inbreeding coefficient was < −0.03, or if none of the following were true: read depth was greater than or equal to 10; genotype quality was greater than or equal to 20; or allele balance was greater than or equal to 0.2. In total, 49,997 exomes were available. Variants were annotated with the Ensembl Variant Effect Predictor version 95 using the --pick-allele flag^72^. LOFTEE was used to identify high-confidence loss of function variants: stop-gain, splice-site disrupting, and frameshift variants^73^.

### Rare variant association test

We conducted a collapsing burden test to assess the impact of loss-of-function variants in 12,168 participants who had both aortic measurements and exome sequencing data available. For quantitative traits (minor axis length of the ascending and descending thoracic aorta), with heritability of approximately 0.6, we estimated that 13 loss-of-function variant carriers would be sufficient to achieve a power of 0.8 at an alpha of 0.05. Variants with MAF >= 0.001 were excluded. Using the LOFTEE “high-confidence” loss-of-function variants, for each of the 3,254 protein-encoding genes with at least 13 carriers of one or more loss-of-function variants in the UK Biobank, we tested whether loss-of-function carrier status was associated with aortic minor axis length. The model was adjusted for weight (kg), height (cm), the MRI serial number, age at enrollment, the cubic natural spline of age at the time of MRI, sex, genotyping array, and PC1-5. We performed an additional analysis that subset the gene list to those within a 500kb radius of one of the independently associated SNPs from the GWAS. These criteria yielded 91 genes (ascending aorta) and 161 genes (descending aorta) for the secondary analysis.

### Association of the ascending aortic polygenic score with incident disease

Within a strictly defined European subset of the UK Biobank, we computed a polygenic score from the 83 autosomal, independently significant SNPs from the ascending aorta GWAS (**Supplementary Table 16**), excluding participants used for the GWAS (**Supplementary Table 17**). We analyzed the relationship between this score and incident thoracic aortic aneurysm or dissection in 391,251 individuals (381 cases) using a Cox proportional hazards model. There is limited data regarding clinical risk factors for thoracic aortic aneurysm outside of associated syndromes and family history, so we chose putatively relevant covariates based in part on inference from evidence in the abdominal aortic aneurysm literature^74^. We adjusted for putative aortic aneurysm risk factors including sex, prevalent diagnoses of type 2 diabetes or hypertension, tobacco smoking history (the number of pack years of tobacco smoking), body mass (the cubic natural spline of BMI), and age (the cubic natural spline of age at enrollment). We also adjusted for other covariates including the cubic natural spline of height, the number of standard alcoholic drinks consumed per week, the genotyping array, and the first five principal components of ancestry.

### Rat aortic nuclei isolation and library generation

Animal experiments were approved by the institutional IACUC at Broad Institute. Wistar rats (Charles River, MA) were acclimated for 2 weeks, with *ad libitum* access to water and chow diet. 17-week-old animals were euthanized between 10am and 12pm using CO2, followed by perfusion with PBS to remove excess blood. Whole aortas - from aortic root to iliac bifurcation - were surgically collected, immediately frozen in LN2 and stored at −80°C until use. For nuclei isolation, aortas were mounted frozen on OCT and sectioned at 60um at −20°C with a cryotome (Leica CM 1950). Nuclei were liberated in ice cold nuclei isolation buffer (NIB: Hepes, Sucrose, MgCl2, KCl, Igepal-630, BSA, pH 7.2) by dounce homogenization. Homogenates were centrifuged at 40g x 4’, at 4°C. Supernatant was filtered through sequential 40um and 10um meshes (Pluriselect, Germany), and filtrate was centrifuged at 600g x 5’, at 4°C. Supernatant was discarded and pellet resuspended and washed once (600g x 5’, 4°C) with nuclei wash buffer (NIB without detergent). Final pellet was resuspended in 150uL nuclei storage buffer (NWB with 1:80 murine RNAse inhibitor, NEB). All procedures were performed on ice. Nuclei, stained with Trypan blue, were manually counted using a hemocytometer (inCyto.com). 7,000 nuclei (5,000 recovery) per aorta were used for droplet generation and library construction according to manufacturer’s protocol (10x Genomics, V2).

### Rat single nucleus RNA sequencing data analysis

Most data analysis was performed using the Terra cloud platform (terra.bio). BCL files for all 9 datasets were processed using *cellranger mkfastq* (CellRanger 3.0.2, 10x Genomics) to generate FASTQ files. These FASTQ files were trimmed using *cutadapt*^75^ to remove the template switch oligo adapter sequence and its reverse complement [AAGCAGTGGTATCAACGCAGAGTACATGGG, CCCATGTACTCTGCGTTGATACCACTGCTT] (max_error_rate=0.07, min_overlap=10) and all four homopolymer repeats [A_30_, C_30_, G_30_, T_30_] (max_error_rate=0.1, min_overlap=20). The trimmed FASTQ files were used as input to *cellranger count* (CellRanger 3.0.2) in order to obtain count matrices.

### Rat transcriptome

The rat transcriptome from Ensembl (*Rattus norvegicus*, Rnor_6.0.96)^76^ lacks full-length Ttn as well as large stretches of other important cardiac-related transcripts including Ryr2. Many other transcripts are annotated with extents shorter than the read alignment would suggest, resulting in low read-mapping to the Ensembl transcriptome. We therefore created an augmented reference transcriptome for the rat which was used for this study.

First, bulk RNA sequencing was generated by strand specific, long insert whole transcriptome sequencing as offered by the Genomics Platform of the Broad Institute (genomics.broadinstitute.org). Briefly, poly-adenylated RNA was isolated from the aorta, AV node, and all four cardiac chambers of two male Wistar rats and converted to sequencing-ready Illumina TruSeq libraries according to manufacturer’s protocols. Libraries were subjected to paired end 50bp sequencing to a mean depth of ∼47,000,000 dually mapping reads per library. A *de novo* reference transcriptome was created from the bulk RNA-seq data using *StringTie* unguided^77^. Only transcripts with at least 5 TPM read evidence were kept. Given that we performed nuclear 3’ scRNA-seq, all transcripts were collapsed to the level of a gene body as we expected to find retained introns in our reads.

Our augmented reference transcriptome was created by starting with Ensembl Rnor_6.0.96, adding Ttn and several other genes from the RGD rat60 reference transcriptome, downloaded from ftp://ftp.rgd.mcw.edu/pub/data_release/GFF3/Gene/Rat/rat60/^78^, and expanding each of the annotations in the Ensembl reference based on two rules: (1) if there is an overlapping gene on the same strand with the same name in RGD, and it does not cause a conflict with another protein-coding Ensembl gene on the same strand, expand the gene definition to match RGD; and (2) if there is an overlapping transcript in the unguided *StringTie* reference, and it does not cause a conflict with any other Ensembl gene on the same strand, expand the gene definition, in whichever direction(s) possible. Compared to the Ensembl Rnor_6.0.96 transcriptome, typically 5-10% more reads from cardiac samples mapped to this amended transcriptome.

### Rat sample-level quality control

Quality control at the level of entire samples was performed by examining QC metrics produced by *cellranger count*, as well as tSNE plots and plots of log(UMI count) versus log(droplet ID) ranked by decreasing UMI count. 3 samples were identified as such strong outliers that they were deemed to be QC failures and subsequently removed.

### Rat snRNA noise removal

Count matrices from the remaining 6 datasets were processed using *cellbender remove-background*^79^ to call cells (and eliminate empty droplets) and to remove background noise caused by ambient RNA and barcode swapping (CellBender 0.1, default settings with expected-cells=5000, total-droplets-included=25000, z-dim=200, z-layers=1000, epochs=300).

### Rat nuclei QC

The number of reads per nucleus mapping to introns, exons, and junctions was tabulated using *scR-Invex* (https://github.com/broadinstitute/scrinvex). Quality control at the level of individual nuclei was performed separately for each sample. QC metrics calculated per nucleus included log(fraction of reads from mitochondrial genes), fraction of reads mapping to exons, and entropy of gene expression. Outlier nuclei were detected using a 3-dimensional Gaussian outlier detection algorithm using the above three QC metrics, fitted on those nuclei with fraction of reads from mitochondrial genes <= 0.05, entropy of gene expression > 6 and < 9, and fraction of reads mapping to exons < 0.35. Outlier detection was performed using the *scikit-learn*^80^ function EllipticEnvelope (contamination=0.02). A distributional cutoff at the 98^th^ percentile of entropy * log (gene count) was used as a surrogate for removing doublets. Between 1500 and 5000 nuclei remained for each sample.

### Rat aorta aggregated map

Count matrices for passing nuclei from each sample were aggregated into one large count matrix. Highly variable genes were computed using Seurat 3^81^ (method=vst, n_genes=2000). Batch effect correction was performed using *scVI*^82^ (latent_dimension=50, max_epochs=150, early_stopping=True, only using highly variable genes). Latent embeddings of each nucleus from scVI were used to create a two-dimensional map using the uniform manifold approximation and projection for dimension reduction (UMAP) algorithm^83^. Nuclei in the aggregated map were clustered using the Louvain algorithm in scanpy^84^, computing nearest-neighbor distances using Euclidean distance in the space of the scVI latent representation. Louvain clustering was run at various resolutions, and the final resolution of 0.8 was chosen manually due to its parsimonious covering of the dataset.

### Differential expression between cell types in rat aorta

Differential expression testing was performed for each gene by comparing expression in a given cluster to all other clusters in R *limma*^85^. Testing was carried out as per the recommendation by Lun and Marioni^86^, after (1) summing count data per sample per cluster, (2) normalizing using DESeq2^87^, and (3) correcting for the mean-variance trend using *voom*. Contrasts of one cell cluster versus all others were fit using the model (∼ 0 + cluster) to extract an estimate of a log fold-change between the given cluster and all others. Only genes where at least two summed sample-clusters showed nonzero expression were tested. Multiple-testing correction was performed using the Benjamini-Hochberg method with a false discovery rate of 0.01. Tens to hundreds of genes were found to be significantly differentially-expressed in each cluster. Cell types were named by examination of the top up-regulated genes in a cluster and manual searching of the literature.

## Data availability

UK Biobank data is made available to researchers from universities and other research institutions with genuine research inquiries, following IRB and UK Biobank approval. Full GWAS summary statistics for ascending and descending thoracic aortic measurements will be available upon publication at the Broad Institute Cardiovascular Disease Knowledge Portal at http://www.broadcvdi.org. Single nucleus RNA sequencing data will be available upon publication at the Single Cell Portal: https://singlecell.broadinstitute.org/single_cell. The dbGAP accession number for aortic phenotypes used in FHS replication is phs000007.v30.p11. All other data are contained within the article and its supplementary information, or are available upon reasonable request to the corresponding author.

## Code availability

The code used to identify connected components is available as a Go library at https://github.com/carbocation/genomisc/tree/master/overlay

## Author Contributions

JPP and PTE conceived of the study. JPP, MDC, SJF, and HL conducted bioinformatic analyses. AA, ADA, and NRT performed the rat lab work. JPP, MEL, and PTE wrote the paper. All other authors contributed to the analysis plan or provided critical revisions.

