## Supplement for "Deep learning enables genetic analysis of the human thoracic aorta"

### Supplementary Material

**Supplementary Material** 1

**Supplementary Note** 1

Association between aortic measurements and other phenotypes 1

**Supplementary Figures** 3

Supplementary Figure 1: Sample Flow Diagram 3

Supplementary Figure 2: Aortic size by age, and sex 4

Supplementary Figure 3: PheWAS with observed aortic traits 5

A) Continuous traits 5

B) Disease phenotypes 5

Supplementary Figure 4: GWAS replication in Framingham 7

Supplementary Figure 5: Genetic correlation with 272 continuous traits 8

Supplementary Figure 6: Genetic correlation between continuous traits and the ascending and descending aorta 9

Supplementary Figure 7: Cell type-specific gene expression at the WWP2 locus 10

Supplementary Figure 8: MAGMA gene set associations 11

**Supplementary References** 12

### Supplementary Note

#### Association between aortic measurements and other phenotypes

Among the participants with aortic imaging data, the size of the thoracic aorta was directly correlated with basal metabolic rate, height, weight, blood pressure, and forced expiratory volume in 1 second, consistent with previous reports[1]. Aortic size was most strongly inversely correlated with heart rate, high-density lipoprotein cholesterol, and sex hormone binding globulin levels (**Supplementary Table 2**; **Supplementary Figure 3A**). We analyzed the association between aortic size and PheCode-based disease labels. As anticipated, the size of the ascending aorta was associated with cardiovascular diseases such as hypertension, aortic aneurysm, and aortic valve disorders, as well as other unexpected traits including varicose veins, obesity, cardiac arrhythmias, cardiomegaly, and osteoarthritis. In contrast, descending thoracic aortic size was associated with obesity, hypertension, and varicose veins, but not with other cardiovascular diseases. In addition, the descending aortic size was directly associated with cholelithiasis and headache, and inversely associated with type 1 diabetes as has previously been observed [2,3](**Supplementary Table 3**; **Supplementary Figure 3B**).

### Supplementary Figures

**Supplementary Figure 1:** Sample Flow Diagram

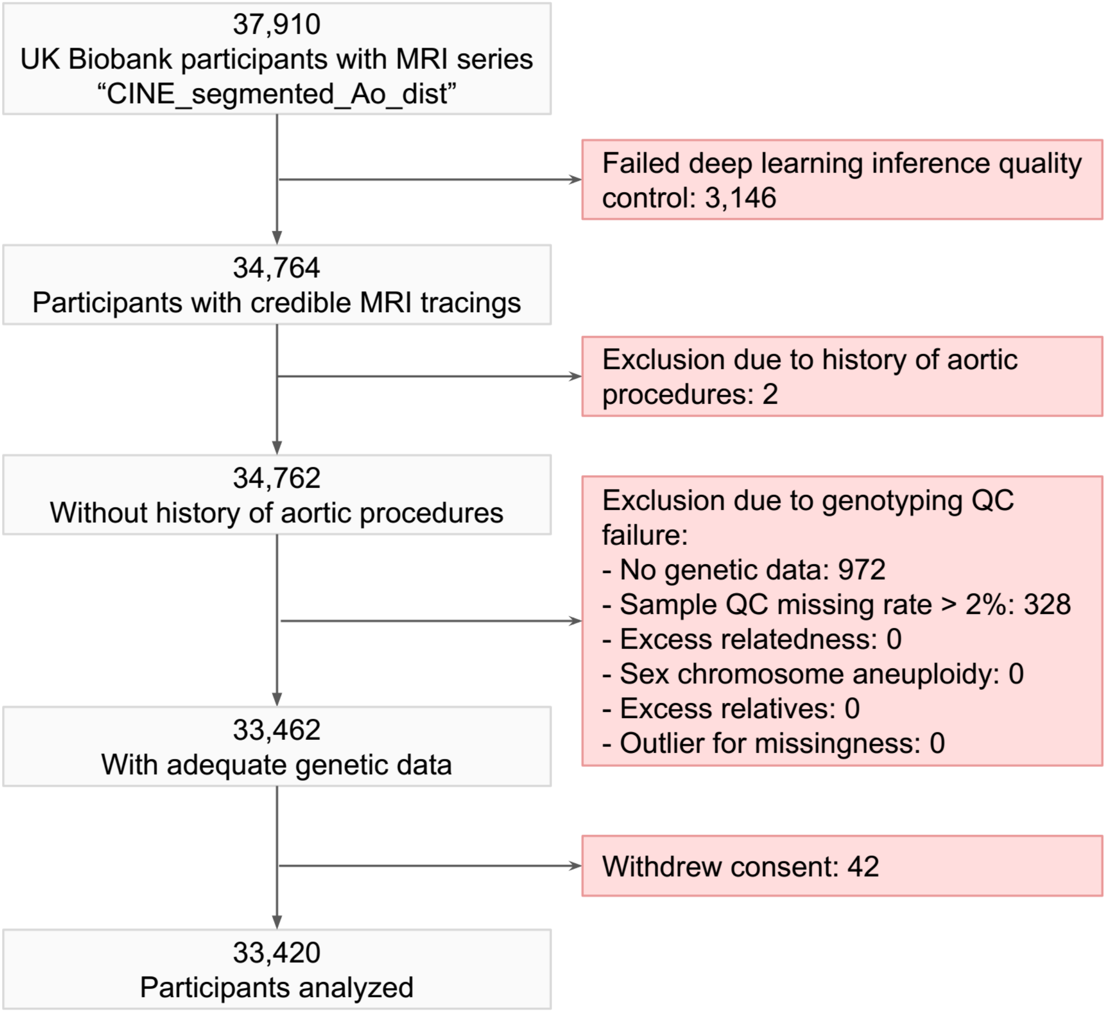

**Supplementary Figure 2:** Aortic size by age, and sex

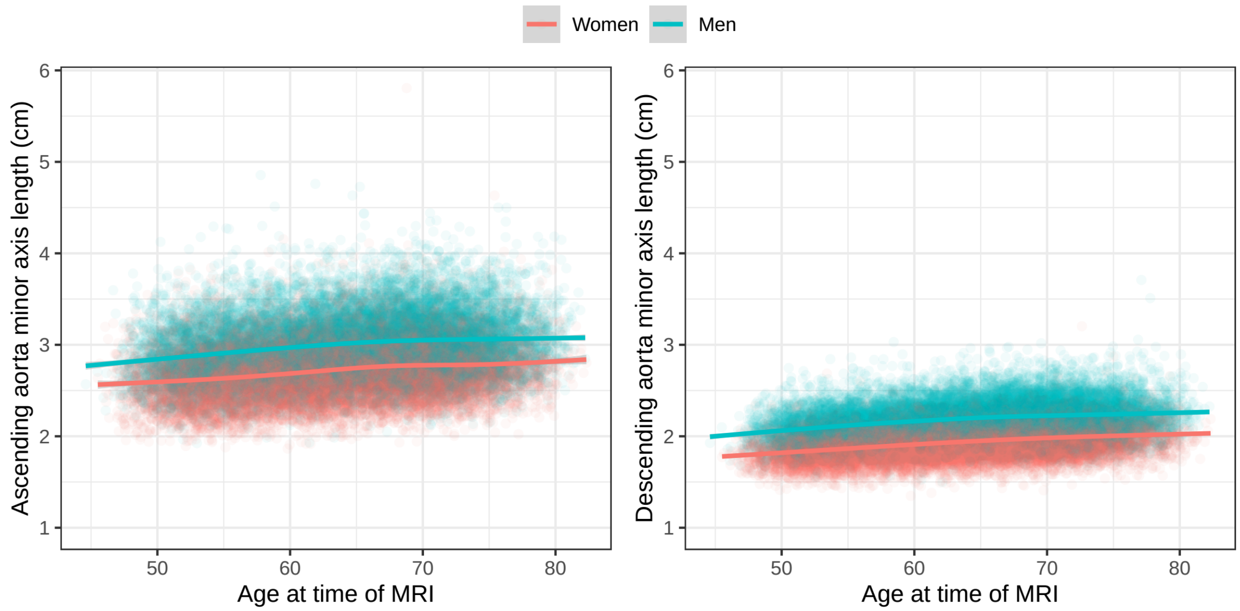


The length of the minor elliptical axis of aorta at its maximum size during the cardiac cycle is shown for the ascending aorta (**left panel**) and the descending aorta (**right panel**). The x axis represents the participant's age at the time of cardiac MRI and the y axis represents the size of aorta. Each point represents one person's measurements; men are plotted in turquoise and women in red. Sex-specific locally weighted scatterplot smoothing (LOESS) curves are overplotted.

**Supplementary Figure 3:** PheWAS with observed aortic traits

A) Continuous traits

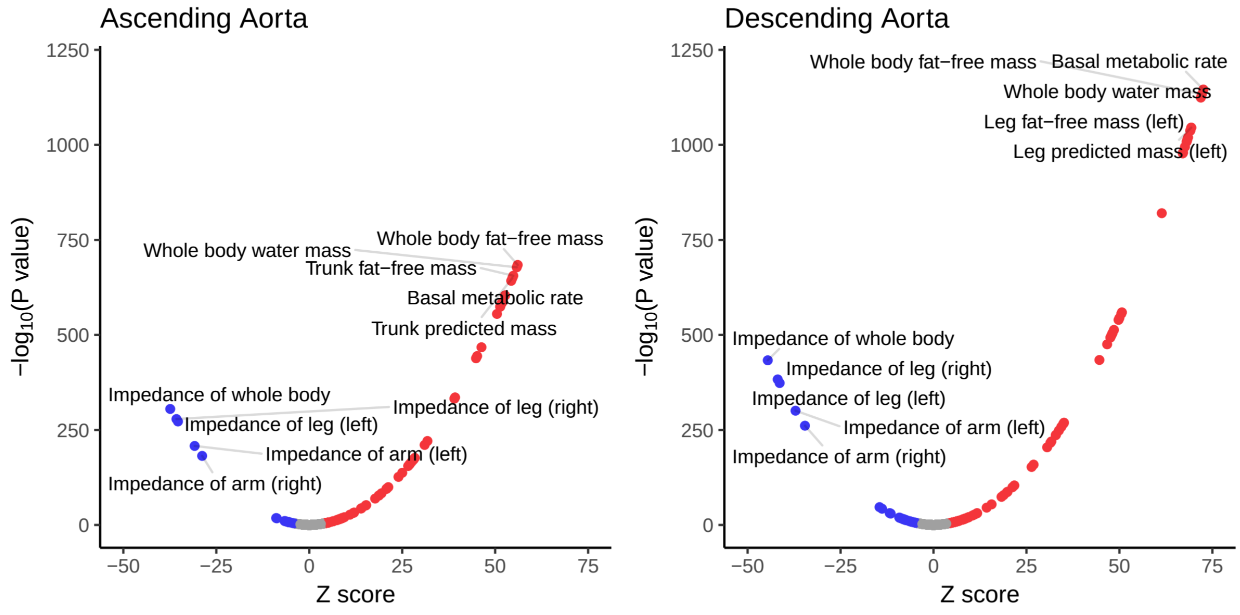


B) Disease phenotypes

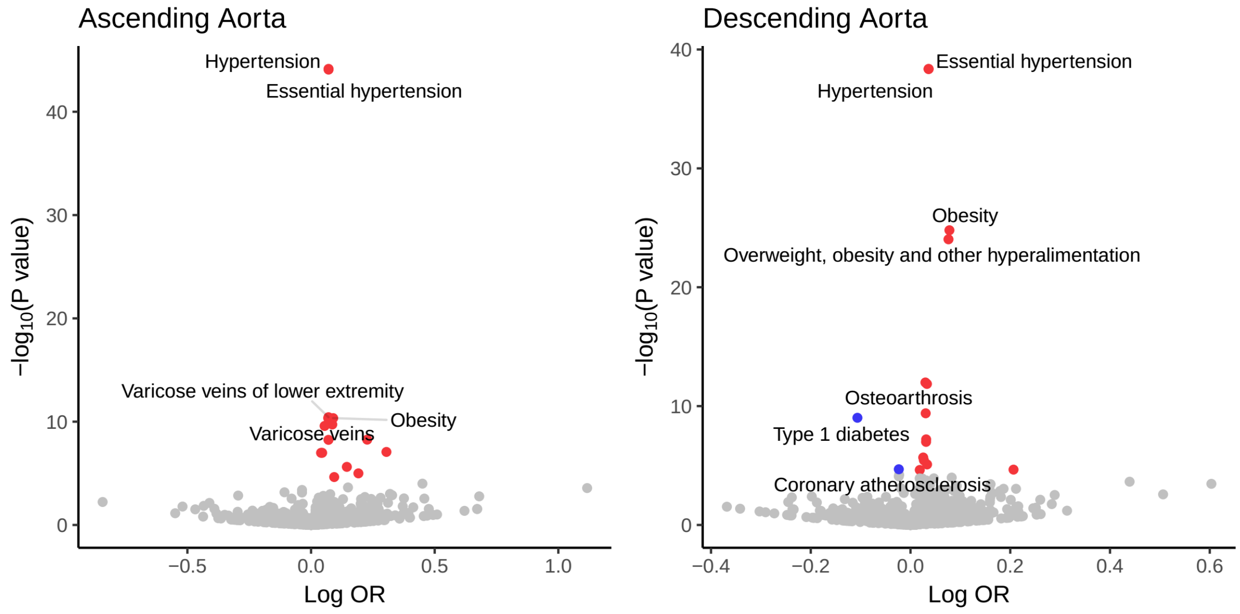


**Panel A**: Phenotypes associated with the size of the ascending (**left panel**) and descending (**right panel**) thoracic aorta are represented in volcano plots. The **x axis** represents the magnitude of the association Z score, while the **y axis** represents the -log10 of the linear model association P value. Traits achieving Bonferroni significance are colored red (positive correlation) or blue (negative correlation). The top 5 positively and negatively correlated traits are labeled. **Panel B**: PheCode-based diseases associated with the size of the ascending (**left panel**) and descending (**right panel**) thoracic aorta. The **x axis** represents the log of the odds ratio for association between disease and aortic size, while the **y axis** represents the -log10 of the association P value in a logistic model. Diseases achieving Bonferroni significance are colored red (positive correlation) or blue (negative correlation). Up to 5 positively and negatively correlated diseases are labeled.

**Supplementary Figure 4:** GWAS replication in Framingham

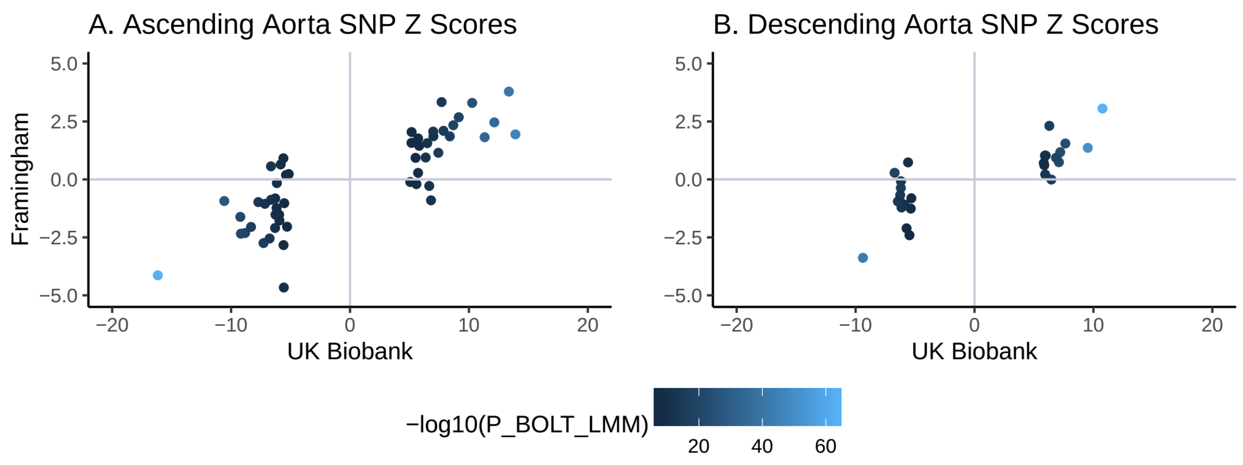


For lead SNPs from the main UK Biobank GWAS that could be identified in a GWAS from FHS, each SNP is plotted based on the UK Biobank Z score (**x axis**) and the FHS Z score (**y axis**). SNPs associated with the ascending aorta are plotted in **Panel A** and those associated with the descending aorta are plotted in **Panel B**. SNP color is based on the -log10 of the association P value from the UK Biobank GWAS.

**Supplementary Figure 5:** Genetic correlation with 272 continuous traits

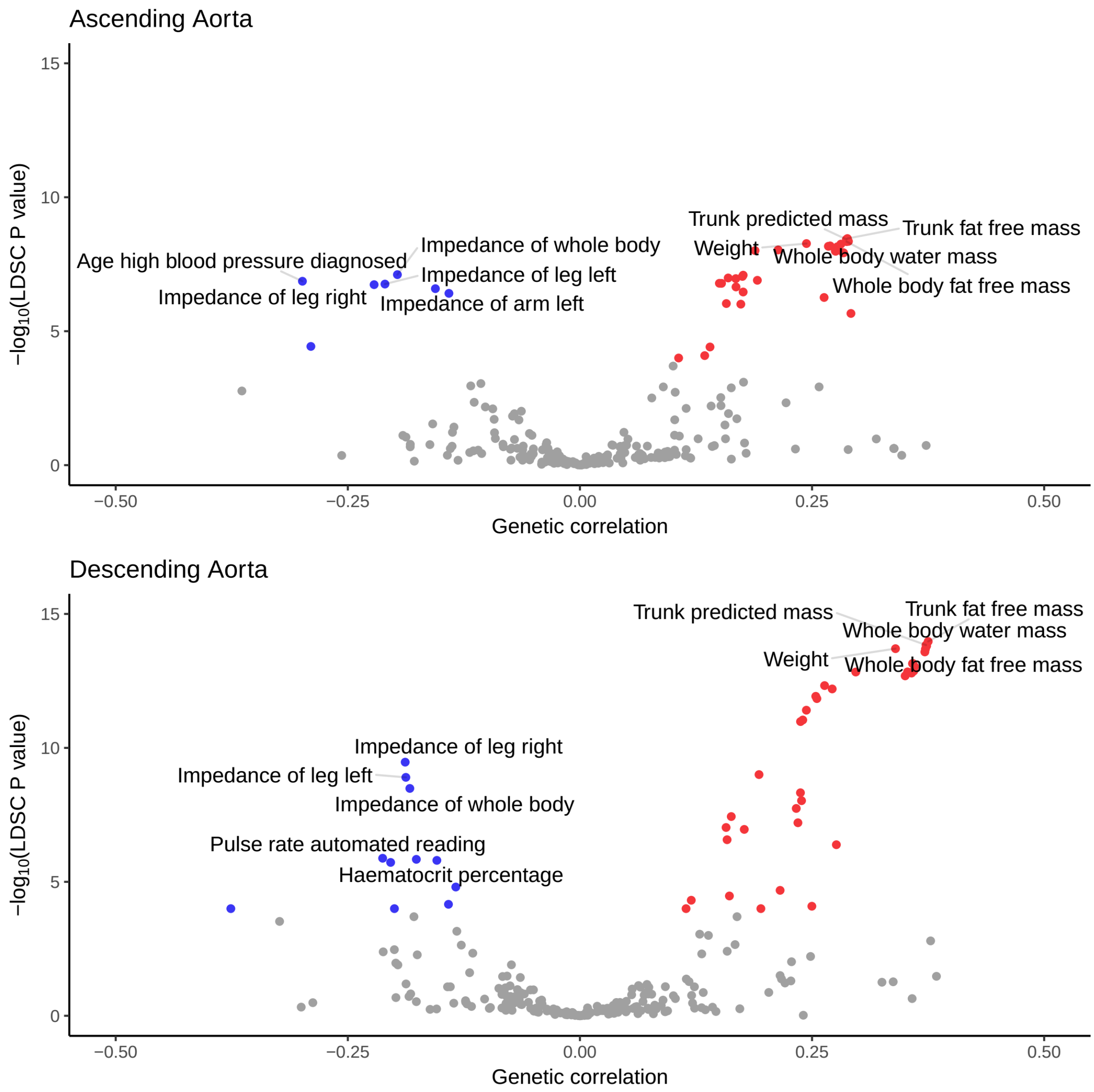


The genetic correlation between 272 continuous traits and the ascending (**top panel**) and descending (**bottom panel**) thoracic aorta are represented in volcano plots. The **x axis** represents the magnitude of genetic correlation, while the **y axis** represents the -log10 of the genetic correlation P value, based on *ldsc*. Traits achieving Bonferroni significance are colored red (for positive genetic correlation) or blue (for negative genetic correlation). The top 5 positively and negatively associated traits are labeled.

**Supplementary Figure 6:** Genetic correlation between continuous traits and the ascending and descending aorta

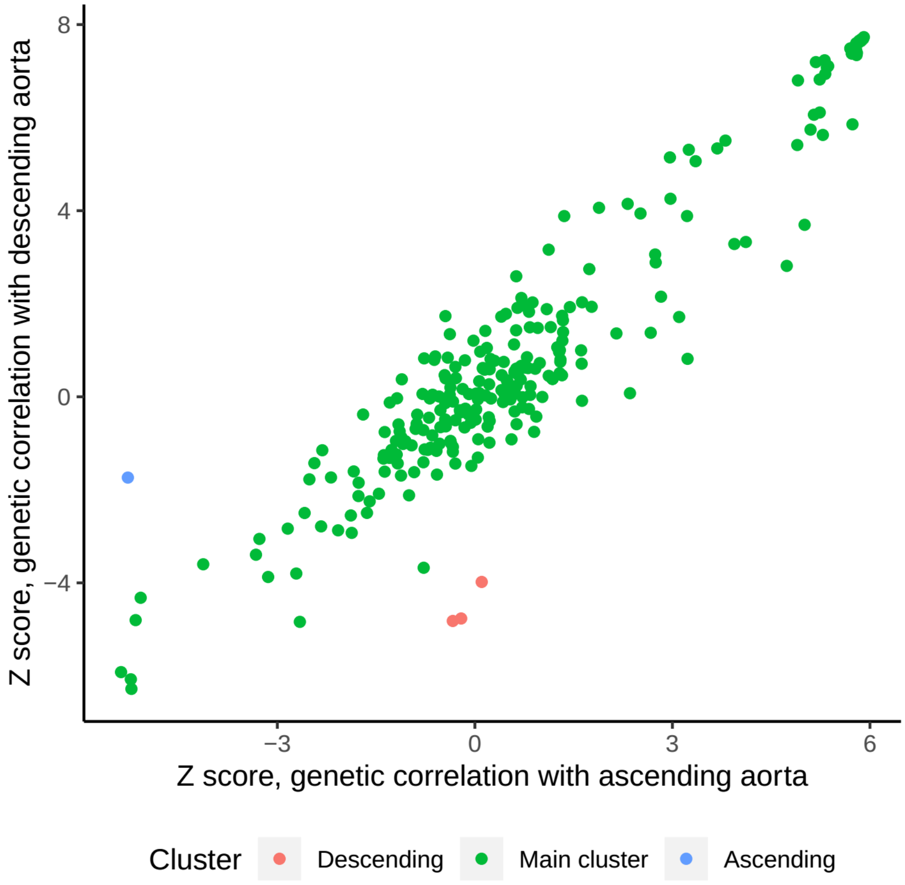


The Z scores for genetic correlation between 272 continuous traits measured by the Neale Lab in the UK Biobank and the ascending and descending aorta are plotted based on association with ascending aorta (**x axis**) and descending aorta (**y axis**). The main inlier group is colored in green. The traits that primarily have a genetic correlation with the ascending aorta are colored in blue (age of onset of hypertension), while the traits that primarily have a genetic correlation with the descending aorta are colored in red (red blood cell counts). The full data is available in **Supplementary Table 6**.

**Supplementary Figure 7**: Cell type-specific gene expression at the *WWP2* locus

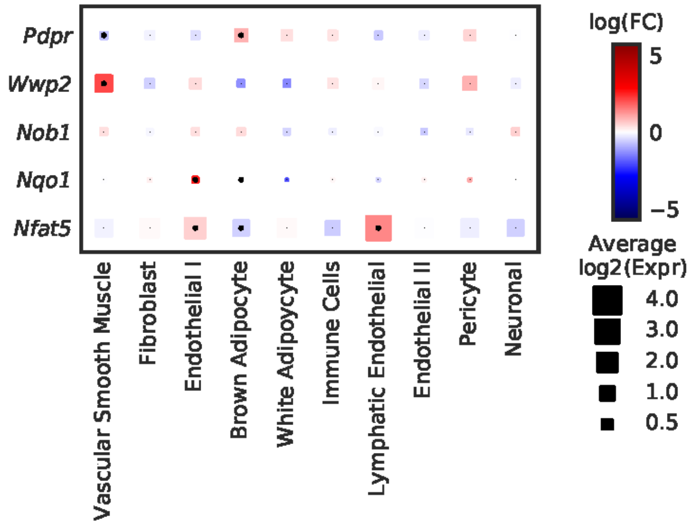


Cell-type specificity of genes with expression data within 500kb of the lead SNP near *WWP2*. As with **Figure 5**, the size of each square represents the average log2(Expr) for a gene across all nuclei in a given cluster. The color represents the log fold-change comparing the expression of the given gene in each cluster to all other clusters based on a formal differential expression model. A dot represents significant up- or down-regulation in the given cluster based on a Benjamini-Hochberg correction for multiple testing at FDR < 0.01. Expr = Normalized nucleus-level expression calculated as the number of counts of a gene divided by the total number of counts in the nucleus and multiplied by 10,000; FC = Fold-change.

**Supplementary Figure 8**: MAGMA gene set associations

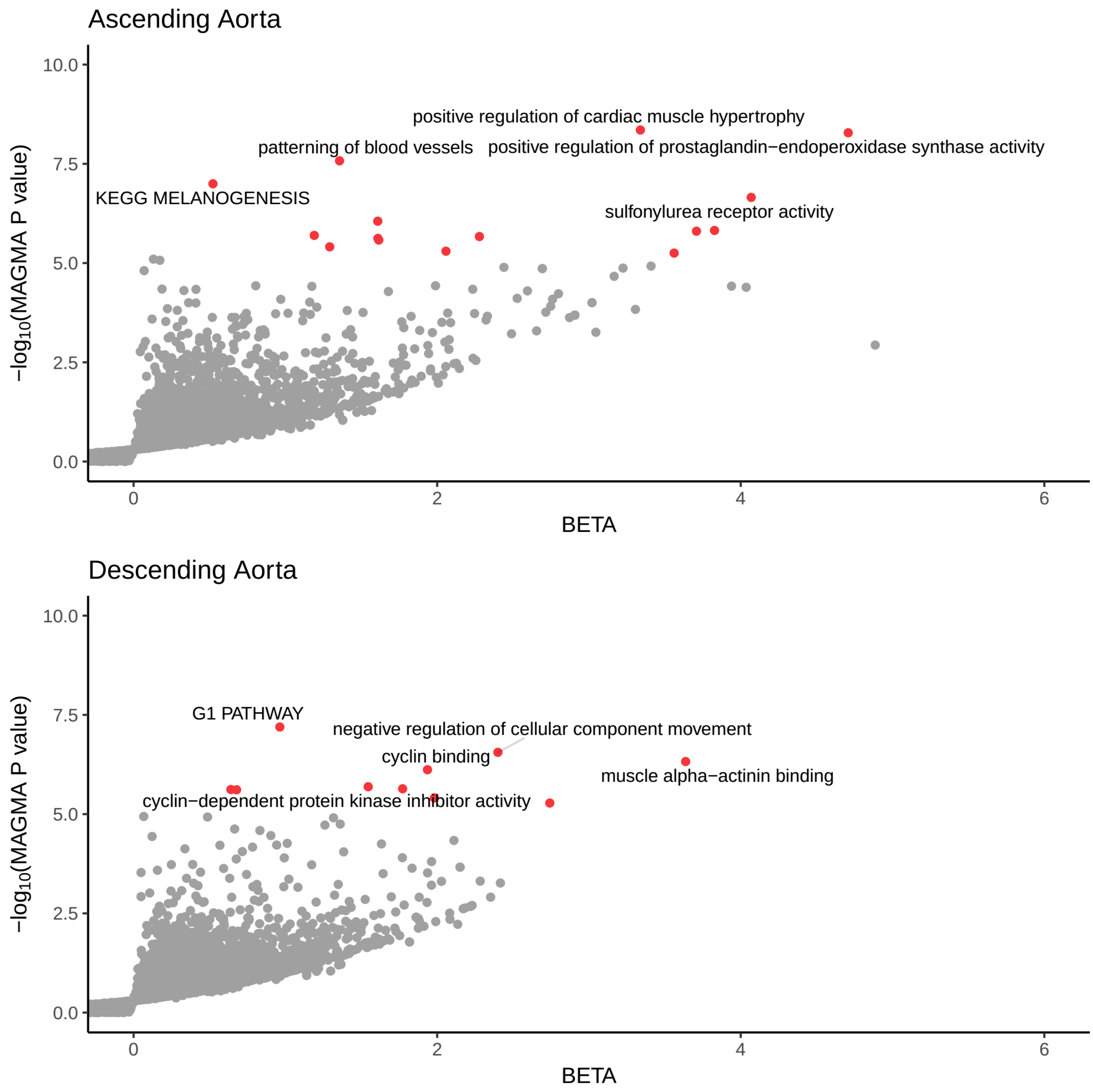


Gene sets enriched in MAGMA analysis of the GWAS of the ascending (**top panel**) and descending (**bottom panel**) thoracic aorta are represented in volcano plots. The **x axis** represents the magnitude of estimated effect of a pathway-based gene set on the aortic trait, while the **y axis** represents the -log10 of the MAGMA association P value. Pathways achieving Bonferroni significance are colored red. The top five pathways are labeled.
